# Crystal and solution structures reveal oligomerization of individual capsid homology domains of Drosophila Arc

**DOI:** 10.1101/2020.11.18.388496

**Authors:** Erik I. Hallin, Sigurbjörn Markússon, Lev Böttger, Andrew E. Torda, Clive R. Bramham, Petri Kursula

**Affiliations:** Department of Biomedicine, University of Bergen, Norway; Centre for Bioinformatics (ZBH), University of Hamburg, Germany; KG Jebsen Centre for Neuropsychiatric Disorders, University of Bergen, Norway; Faculty of Biochemistry and Molecular Medicine & Biocenter Oulu, University of Oulu, Finland

**Author notes:** equal contribution (Erik I. Hallin, Sigurbjörn Markússon).

## Abstract

Synaptic plasticity is vital for brain function and memory formation. One of the key proteins in long-term synaptic plasticity and memory is the activity-regulated cytoskeleton-associated protein (Arc). Mammalian Arc forms virus-like capsid-like structures in a process requiring the N-terminal domain and contains two C-terminal lobes that are structural homologues to retroviral capsids. *Drosophila* has two isoforms of Arc, dArc1 and dArc2, with low sequence similarity to mammalian Arc, but lacking the mammalian Arc N-terminal domain. Both dArc isoforms have a capsid homology domain consisting of N- and C-terminal lobes. We carried out structural characterization of the four individual dArc lobe domains. As opposed to the corresponding mammalian Arc lobe domains, which are monomeric, the dArc lobes were all oligomeric in solution, indicating a strong propensity for homophilic interactions. The N-lobe from dArc2 formed a domain-swapped dimer in the crystal structure, resulting in a novel dimer interaction that could be relevant for capsid assembly or other dArc functions. This domain-swapped structure resembles the dimeric protein C of flavivirus capsids, as well as the structure of histones dimers, domain-swapped transcription factors, and membrane-interacting BAK domains. The strong oligomerization properties of the isolated dArc lobe domains explain the ability of dArc to form capsids in the absence of any large N-terminal domain, in contrast to the mammalian protein.

## Introduction

Memory formation in the brain is dependent on synaptic plasticity, and the activity-regulated cytoskeleton-associated protein (Arc) plays an important role in this process [1,2]. Arc promotes the endocytosis of AMPA receptors located on the post-synaptic membrane [3–5], regulates actin cytoskeletal dynamics and dendritic spine structure [6–8], and enters the nucleus to regulate gene expression [4,9,10]. The targeting of AMPA receptors may involve direct interactions of stargazin (TARPγ2) with both AMPA receptors and Arc [11–13]. Due to its many interaction partners, Arc regulates several neuronal signalling processes as well as the structure of the postsynaptic density scaffold [2,14].

Arc forms capsid-like structures that may transfer information from one neuron to another [15,16]. Mammalian Arc (mArc) has a C-terminal domain (Arc-CT) with close structural homology to the C-terminal domain (CA-CTD) of the retroviral capsid (CA) protein [12], and mArc-CT consists of two structurally similar lobe domains, N-lobe (NL) and C-lobe (CL) [12]. Viral CA has in addition an N-terminal domain (CA-NTD), and both CA-NTD and CA-CTD are involved in viral capsid assembly. mArc has a large N-terminal domain (Arc-NT) of unknown structure, which is absent in dArc. The Arc-NT is predicted to have homology to the retroviral matrix domain and is required for the formation of large mArc oligomers. Without its N-terminal domain, mArc is monomeric in solution [17]. In mArc, it is likely that the presence of both mArc-NT and mArc-CT are required for high-order oligomerization and capsid formation [18,19].

*Drosophila* has two Arc isoforms (dArc1 and dArc2), which share high sequence similarity. Drosophila Arc (dArc) isoforms have a CT domain, containing tandem N-and C-lobes, but lack an Arc-NT found in mArc. However, dArc is able to form capsids [16], whose structure has been determined by electron cryomicroscopy [20]. Whether dArc functions similarly to mArc in neurons, even if the functionally important mArc-NT is missing and the sequence similarity to mArc is low, is currently unknown. Mammalian Arc also forms capsids [16], but the high-resolution structure remains to be solved.

We solved crystal structures of the individual dArc lobe domains. The CL of both dArc1 and dArc2 is structurally homologous to the mArc lobe domains, confirming the connection to mArc and retroviral capsids. The structure of dArc2-NL showed a domain-swapped dimer, resulting in a structure similar to the flavivirus capsid protein and resembling histones as well as membrane-interacting BAK domains. All individual dArc lobes are oligomeric in solution, in contrast to the monomeric mArc-CT. Such oligomeric units could be building blocks during the assembly of virus-like capsids by full-length dArc, or they could relate to its other functions.

## Materials and methods

### Recombinant protein production

Proteins were expressed in Escherichia coli BL21(DE3) with a TEV protease-cleavable His tag-maltose binding protein (MBP) fusion at the N terminus. Cells were grown at +37 °C until an A_600_ of 0.6 was reached. 1 mM isopropyl β-D-1-thiogalactopyranoside was added to start the induction, lasting 4 h at +30 °C. The cells were lysed in HBS (40 mM Hepes, pH 7.5, 100 mM NaCl) containing 0.1 mg/ml lysozyme, by one freeze-thaw cycle followed by sonication. The lysate was centrifuged at 16 000 g for 30 min at +4 °C and loaded onto a Ni-NTA resin. After washing with HBS containing 20 mM imidazole, the protein was eluted with HBS containing 300 mM imidazole. His-tagged TEV protease [21] was added to the eluate, and the sample was dialyzed against 20 mM Hepes (pH 7.5), 100 mM NaCl, and 1 mM dithiothreitol for 20 h at +4 °C. The sample was passed through a Ni-NTA resin again to remove the TEV protease and the cleaved His-MBP tag.

For the purification of dArc1-CL and dArc2-CL, an additional step was required to remove remains of the cleaved MBP tag. This was done by passing the sample through an amylose resin, equilibrated with HBS containing 1 mM EDTA.

The Ni-NTA or amylose flow-through was loaded on a Superdex S200 16/600 column, equilibrated with TBS (20 mM Tris-HCl (pH 7.4), 150 mM NaCl). All proteins gave one major peak in the chromatogram. Selected fractions were concentrated using spin concentrators to a final concentration of 10 mg/ml. Protein purity was analyzed using sodium dodecyl sulphate–polyacrylamide gel electrophoresis, giving one strong Coomassie-stained band of the expected size. Protein identity was confirmed using mass spectrometry of trypsin-digested in-gel samples, as described [22].

The details of the protein constructs are given in S1 Table. The expression and purification of human Arc NL and CL have been described [17].

### Size exclusion chromatography – multi-angle light scattering

The absolute mass of the proteins was determined by SEC-MALS, using a miniDawn Treos instrument (Wyatt). A Superdex S200 Increase 10/300 equilibrated with TBS was used for sample separation. The system was calibrated using bovine serum albumin, and the protein concentration was measured using an online refractometer. Data were analysed with ASTRA (Wyatt).

### Circular dichroism spectroscopy

The ellipticity of the proteins was recorded using a Jasco J-810 spectropolarimeter and a 1-mm quartz cuvette. The protein concentration was 0.2 mg/ml in 20 mM phosphate (pH 7). The experiments were done at +20 °C.

### X-ray crystallography and structure analysis

Crystals were obtained by sitting-drop vapor diffusion at +20 °C. The crystals of dArc2-NL were grown by mixing 150 nl of dArc2-NL at 8 mg/ml with 150 nl of reservoir solution (200 mM ammonium chloride, 100 mM sodium acetate (pH 5), 20 % PEG 6000). The crystals of dArc2-CL were made by mixing 200 nl of protein at 15 mg/ml with 100 nl of reservoir solution (1.25 M ammonium sulphate, 100 mM Tris (pH 8.5), 200 mM lithium sulphate). The crystals of dArc1-CL were made by mixing 150 nl of the protein at 12 mg/ml with 150 nl of a reservoir, consisting of 100 mM MIB buffer (malonic acid, imidazole, boric acid) (pH 5) and 25% PEG 1500. The crystals of dArc1-CL used for phasing were grown by mixing 2 μl of the purified protein at 12 mg/ml with 2 μl of a reservoir solution, consisting of 20% PEG 3350, by hanging-drop vapour diffusion at +20 °C. These crystals were soaked in a solution of 20% PEG 3350 with 500 mM NaI for 20 s.

Crystals were mounted in loops and snap-frozen in liquid nitrogen. X-ray diffraction data for dArc2-NL were collected on the I03 beamline at Diamond Light Source (Oxfordshire, UK), while the data for dArc1-CL and dArc2-CL were collected on the P11 beamline [23] at PETRAIII/DESY (Hamburg, Germany). All data were processed using XDS [24].

Phasing of dArc2-NL was done with molecular replacement in AMPLE [25] and ab initio models generated by QUARK [26], on the CCP4 online server [27,28]. The phasing of dArc1-CL was done using iodine single-wavelength anomalous dispersion (SAD) and the Auto-Rickshaw pipeline [29], with the combined use of SHELX [30], PHASER [31], PARROT [32], and BUCCANEER [33]. The resulting near-complete model was taken as a template for molecular replacement in PHASER [31], using atomic-resolution data from a native crystal with a different space group. The phasing of dArc2-CL was done using the dimeric structure of dArc1-CL as a search model in PHASER [31]. All structures were refined with phenix.refine [34], and model building was done in Coot [35]. The quality of the structures was assessed using MolProbity [36]. Data processing and refinement statistics are given in Table 1.

**Table 1.**
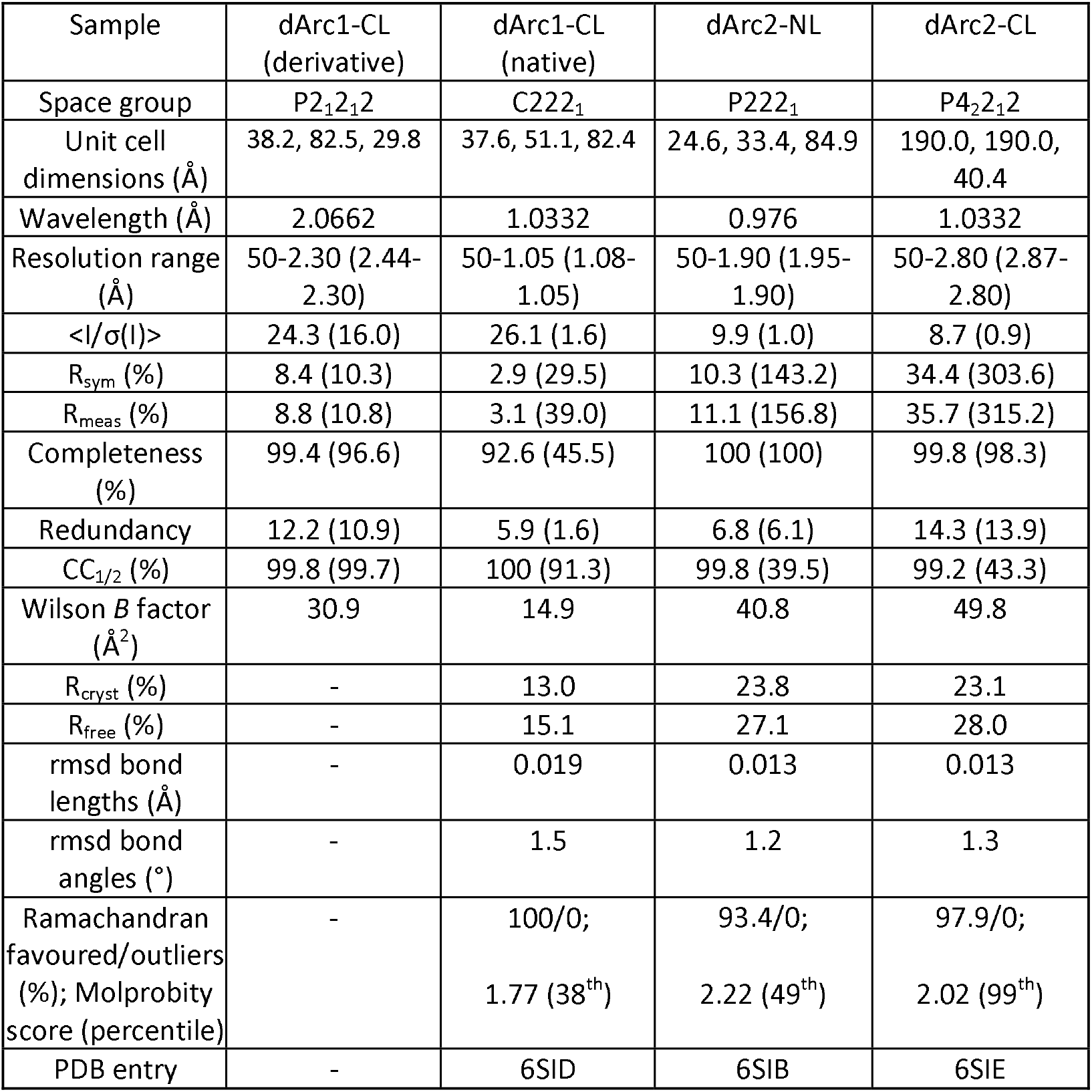
Data processing and structure refinement. The values in parentheses refer to the highest-resolution shell.

| Sample | dArc1-CL<br>(derivative) | dArc1-CL<br>(native) | dArc2-NL | dArc2-CL |
| --- | --- | --- | --- | --- |
| Space group | P2<sub>1</sub>2<sub>1</sub>2 | C222<sub>1</sub> | P222<sub>1</sub> | P4<sub>2</sub>2<sub>1</sub>2 |
| Unit cell<br>dimensions (Å) | 38.2, 82.5, 29.8 | 37.6, 51.1, 82.4 | 24.6, 33.4, 84.9 | 190.0, 190.0,<br>40.4 |
| Wavelength (Å) | 2.0662 | 1.0332 | 0.976 | 1.0332 |
| Resolution range<br>(Å) | 50-2.30 (2.44-2.30) | 50-1.05 (1.08-1.05) | 50-1.90 (1.95-1.90) | 50-2.80 (2.87-2.80) |
| $\langle I/\sigma(I) \rangle$ | 24.3 (16.0) | 26.1 (1.6) | 9.9 (1.0) | 8.7 (0.9) |
| R<sub>sym</sub> (%) | 8.4 (10.3) | 2.9 (29.5) | 10.3 (143.2) | 34.4 (303.6) |
| R<sub>meas</sub> (%) | 8.8 (10.8) | 3.1 (39.0) | 11.1 (156.8) | 35.7 (315.2) |
| Completeness<br>(%) | 99.4 (96.6) | 92.6 (45.5) | 100 (100) | 99.8 (98.3) |
| Redundancy | 12.2 (10.9) | 5.9 (1.6) | 6.8 (6.1) | 14.3 (13.9) |
| CC<sub>1/2</sub> (%) | 99.8 (99.7) | 100 (91.3) | 99.8 (39.5) | 99.2 (43.3) |
| Wilson B factor<br>(Å<sup>2</sup>) | 30.9 | 14.9 | 40.8 | 49.8 |
| R<sub>cryst</sub> (%) | - | 13.0 | 23.8 | 23.1 |
| R<sub>free</sub> (%) | - | 15.1 | 27.1 | 28.0 |
| rmsd bond<br>lengths (Å) | - | 0.019 | 0.013 | 0.013 |
| rmsd bond<br>angles (°) | - | 1.5 | 1.2 | 1.3 |
| Ramachandran<br>favoured/outliers<br>(%); Molprobability<br>score (percentile) | - | 100/0;<br>1.77 (38<sup>th</sup>) | 93.4/0;<br>2.22 (49<sup>th</sup>) | 97.9/0;<br>2.02 (99<sup>th</sup>) |
| PDB entry | - | 6SID | 6SIB | 6SIE |

### Structure analysis

The PISA server [37] was used to calculate probable oligomeric states from crystal symmetry, and PDBsum [38] and PISA were used for structural analysis and in-depth analysis of dimer interfaces. Structural homologues were searched using DALI [39] and SALAMI [40], in addition to the known homologues from literature and manual searches. Electrostatic potential maps were calculated with PDB2PQR and APBS [41] and visualised in UCSF Chimera [42] or PyMOL. Sequence identity between the dArc CA lobes and structural homologues was calculated using the EMBOSS needle server [43], and the dArc1-NL homology model was generated using the SWISS-MODEL server [44], with the dArc2-NL crystal structure as template.

### Small-angle X-ray scattering

Synchrotron SAXS data for dArc1-NL and dArc2-NL were collected on the B21 beamline at Diamond Light Source (Oxfordshire, UK) using a SEC-SAXS setup, where SAXS frames are collected as the sample elutes from a SEC column. A Superdex S200 Increase 3.2 column equilibrated with TBS was used. The injected sample was at 5 mg/ml, and the measurements were done at +10 °C. SEC-SAXS data for dArc1-CL and dArc2-CL were similarly collected on the P12 beamline [45] at EMBL/DESY (Hamburg, Germany).

SAXS data were processed using ATSAS [46], and the frames showed no signs of aggregation or radiation damage. Ab initio dummy atom and chain-like SAXS models were built with DAMMIN [47] and GASBOR [48], respectively. CRYSOL [49] was used to generate SAXS scattering profiles from 3D protein structures and compare these to experimental SAXS data.

### Isothermal titration calorimetry

A MicroCal iTC200 instrument (Malvern, UK) was used to determine the binding affinity of a stargazin peptide (RIPSYRYR with N-terminal acetylation and C-terminal amidation) to the NL of *Drosophila* and human Arc. Arc in the cell had a concentration either 0.5 mM (dArc N-lobes) or 0.25 mM (hArc N-lobe); the peptide concentration in the syringe was 10-fold higher. The peptide was injected in 26 3-μl injections, with an initial injection of 0.5 μl and a second 0.5-μl injection after 14 injections due to syringe refill. Both the protein and peptide were in TBS buffer. The experiments were done at +25 °C, and the data were analyzed with MicroCal Origin 7, using a one-site binding model.

### Sequence comparisons

For conservation calculations, homologues from the non-redundant sequence database were collected using BLAST, accepting hits with an *e*-value less than 10^−10^ [50]. For larger sets and phylogenetic speculation, homologues were collected using iterative PSI-BLAST in up to three stages, each with no more than four iterations, accepting homologues with *e*-values less than 10^−20^, 10^−10^, and 10^−8^[51]. Full length proteins or the candidate ranges were re-aligned using MAFFT in its most accurate mode, with up to 200 iterations [52]. Before alignment, redundancy amongst the sequences was removed by calculating an alignment in fast mode, saving the matrix of distances between sequences, and sorting the list of pair distances. Starting from the smallest distance, one member of each pair was removed until the target number was reached. This removes redundancy and ensures the most even spread of sequences within a set of homologues. For conservation calculations, search starting points were NP_610955.1 (dArc1-NL and full-length dArc1), as well as PDB codes 6sib for dArc2-NL, 6sid for dArc1-CL, and 6sie for dArc2-CL.

Sequence conservation/variability was calculated from the alignments using entropy,

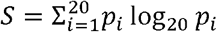

where *p_i_* is the frequency of amino acid type *i* at a given alignment position.

## Results and discussion

Although mArc and dArc both contain similar lobe domains (Fig 1), In mArc, low solubility is caused by the NT [17]. A construct containing both the NL and CL is soluble and fully monomeric for hArc [17]. dArcs lack an Arc-NT, but they are predicted to have an N-terminal helix that might play a role in oligomerization. The isolated NL and CL domains of both dArc1 and dArc2 could be produced in soluble form, while dArc constructs containing both lobes were insoluble (data not shown); this low solubility resembles that of full-length mArc. The different behaviour of the isolated lobe domains suggests that mArc and dArc differ in the mechanism, with which they form larger structures such as capsids. Our aim was to understand the structural basis of these differences.

**Fig 1.**
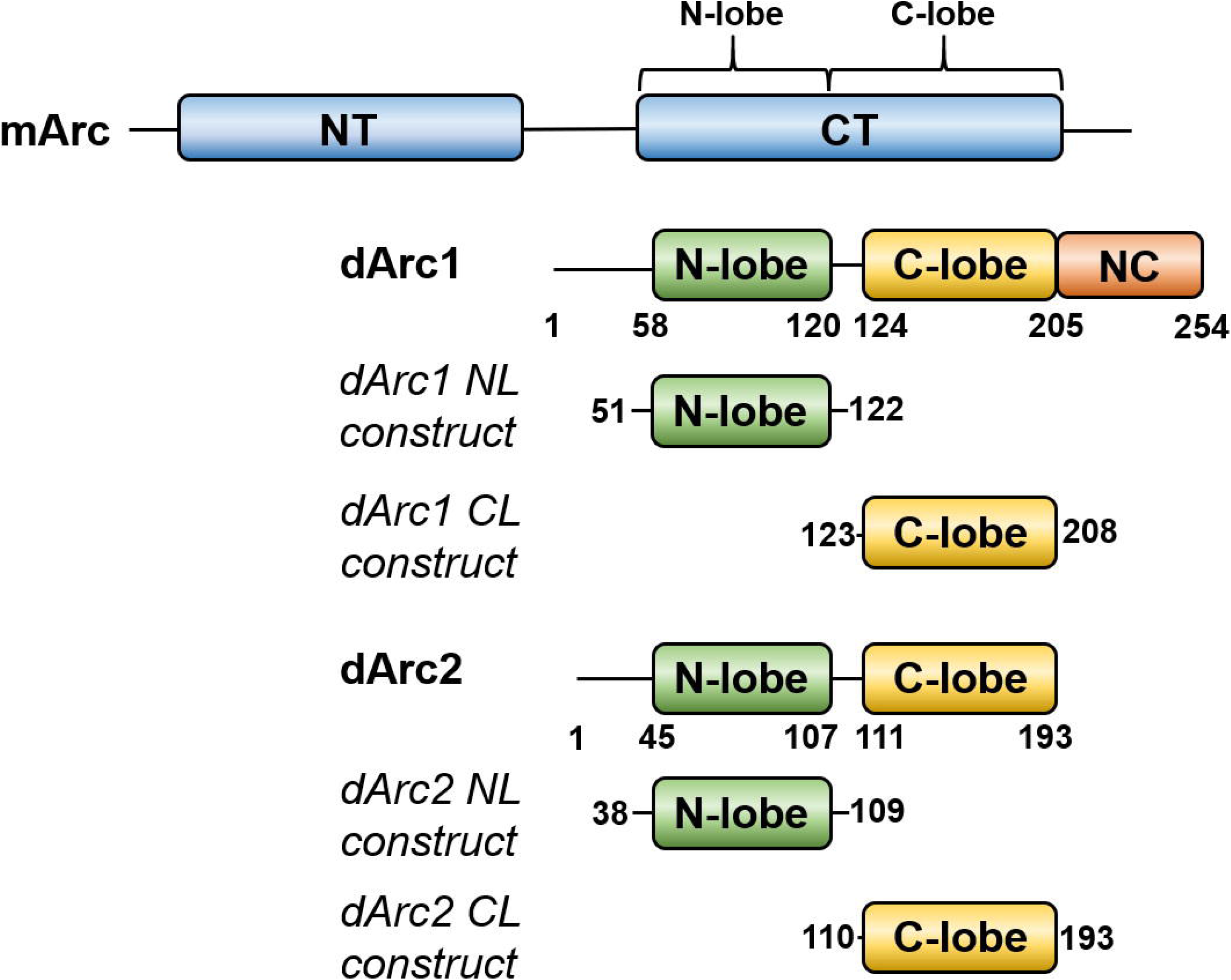
Comparison of mArc and dArc domain structure. The constructs used for structural studies on the individual dArc lobe domains are indicated.

### Crystal structure of dArc2-NL presents a domain-swapped dimer similar to nucleotide- and membrane-interacting proteins

It has been suggested that viral CA-CTD domains employ different dimerization modes during capsid assembly [53]. In the crystal, dArc2-NL is a domain-swapped dimer, in which the second and third helices of the canonical lobe domain are fused, forming an extended helix (Fig 2A-C). One layer of the dimer is formed by α2 of each subunit, which pack at a cross angle of 140.7°. The other layer is formed by the remaining helices of the α2 bundle, and α1 lies in a groove formed between α2 and α3. The subunit interface of the dimer spans 3440 Å^2^ of buried surface area. The interface consists exclusively π-π and other van der Waals interactions, and the helices encapsulate a hydrophobic core between the monomers, most prominently by Phe51, Val55, Pro74, Phe77, Ile80, Trp84, Trp95, Leu99, Leu102, and Phe106 (Fig 2B). The solvent-exposed surface of the dimer is charged and polarised (Fig 2D); the electrostatic surface potential of the side formed by the two coiled α2 helices is highly positive, while the opposing surface formed by α1 and α3 is mainly negative. In the extended crystal lattice (Fig S1), these opposing charges take part in crystal contacts. The crystal packing does not, however, explain the tetrameric oligomerization state of the protein in solution (see below).

**Fig 2.**
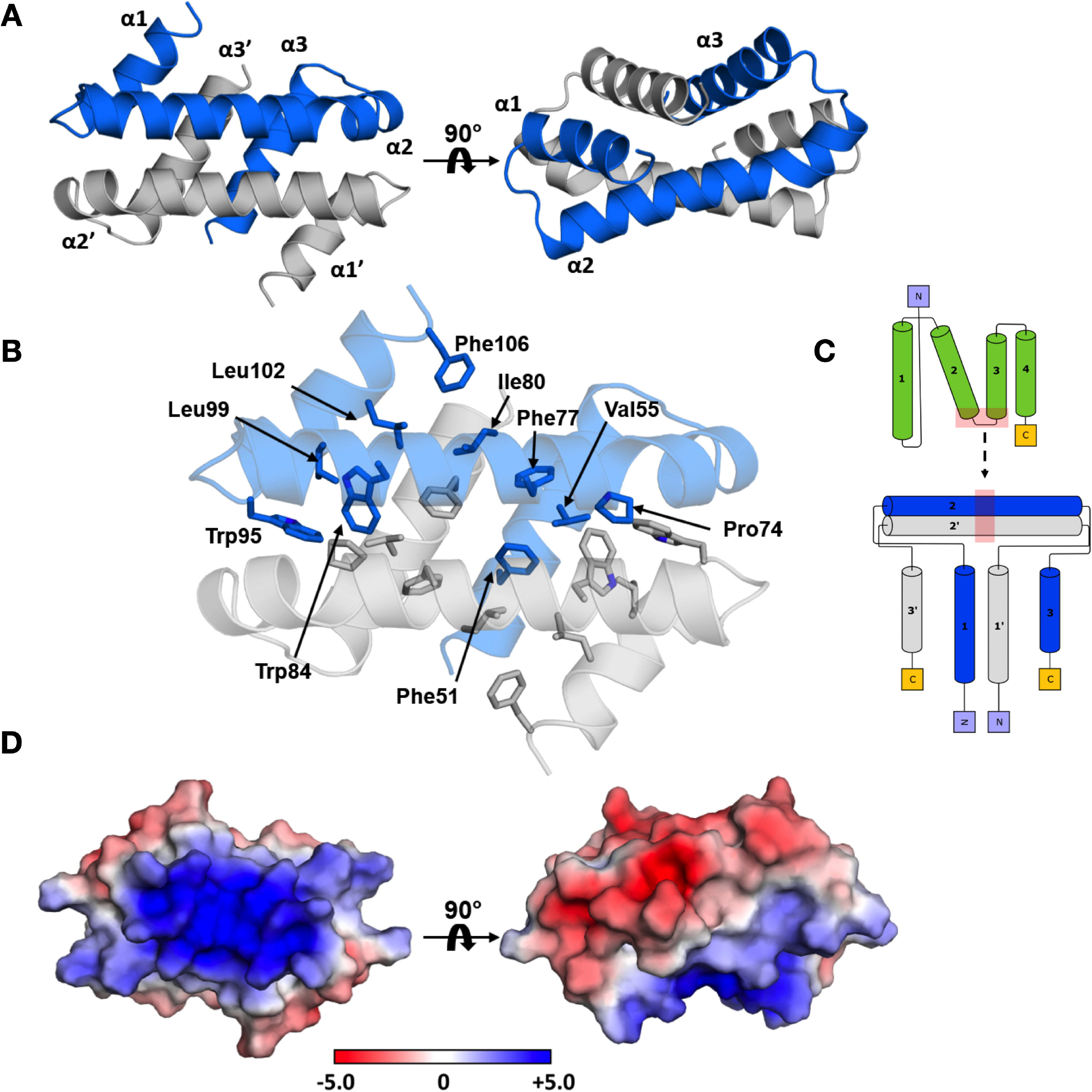
The crystal structure of dArc2-NL. (A) The domain swapped dimer observed in the crystal, with the three α-helices labeled. (B) The folded dimer encapsulates an extensive hydrophobic core, with no polar interactions connecting the two dimers. Residues are only labelled in subunit A, but also seen in subunit B. (C) Topology diagram of domain swapping. (D) The electrostatic surface of the dimer, calculated using APBS. The longitudinal axes of α2 and α2’ have a positive surface potential, in contrast to the surface formed by α1, α3, α1’, and α3’. The protein is in the same orientations as in (A).

The dArc2-NL domain-swapped dimer differs from the dimers of retroviral capsid CA-CTD seen with crystallography and NMR [54] and might be functionally relevant for dArc oligomerization and capsid formation. However, the cryo-EM structures of the dArc1 and dArc2 capsids [20] do not show such domain swapping, and the dArc-NL forms penta- and hexameric rings instead (Fig 3A). In both capsids, a kink is introduced in the extended α2 (at Leu76, Phe77, and Lys78 in dArc2), resulting in breaking of the helix into two to form a canonical lobe domain fold. As a result, the surface charges of the lobe are re-oriented such that the negatively charged surface of α1 can interact with α2 (Fig 3A). Therefore, the turn seems vital to the formation of the capsid hexa- and pentamers. An interesting discrepancy between the domain-swapped dimer, the capsid structure, and the recently published crystal structure of the dArc1 CT domain [55], is the presence of the N-terminal tail preceding the NL (residues 29-44 in dArc2, residues 41-57 in dArc1). In the capsid structures of both dArc isoforms, the tail packs into the exposed hydrophobic core of the lobe (Fig 3B), and Phe32 and Phe39 are observed in two hydrophobic pockets and Ser40 interacts directly with Lys78 and Ser79, of the α2 kink, via hydrogen bonding. A part of the tail (residues 37-45) was included in the dArc2-NL crystallized here. However, this fragment could not be resolved, presumably due to flexibility. Therefore, it seems that the full N-terminal fragment, including Phe32, is needed for the packing of the tail into the exposed hydrophobic core of the lobe and the formation of the penta- and hexameric forms found in capsids. The binding surface for the N-terminal segment is buried at the interface of the dArc2-NL domain-swapped dimer (Fig 2A-B). The structure of dArc2-NL reveals an intrinsic property of the Arc lobe domain to form alternate dimers via domain swapping, possibly regulated through interactions of the folded dArc2-NL core domain with the N-terminal tail.

**Fig 3.**
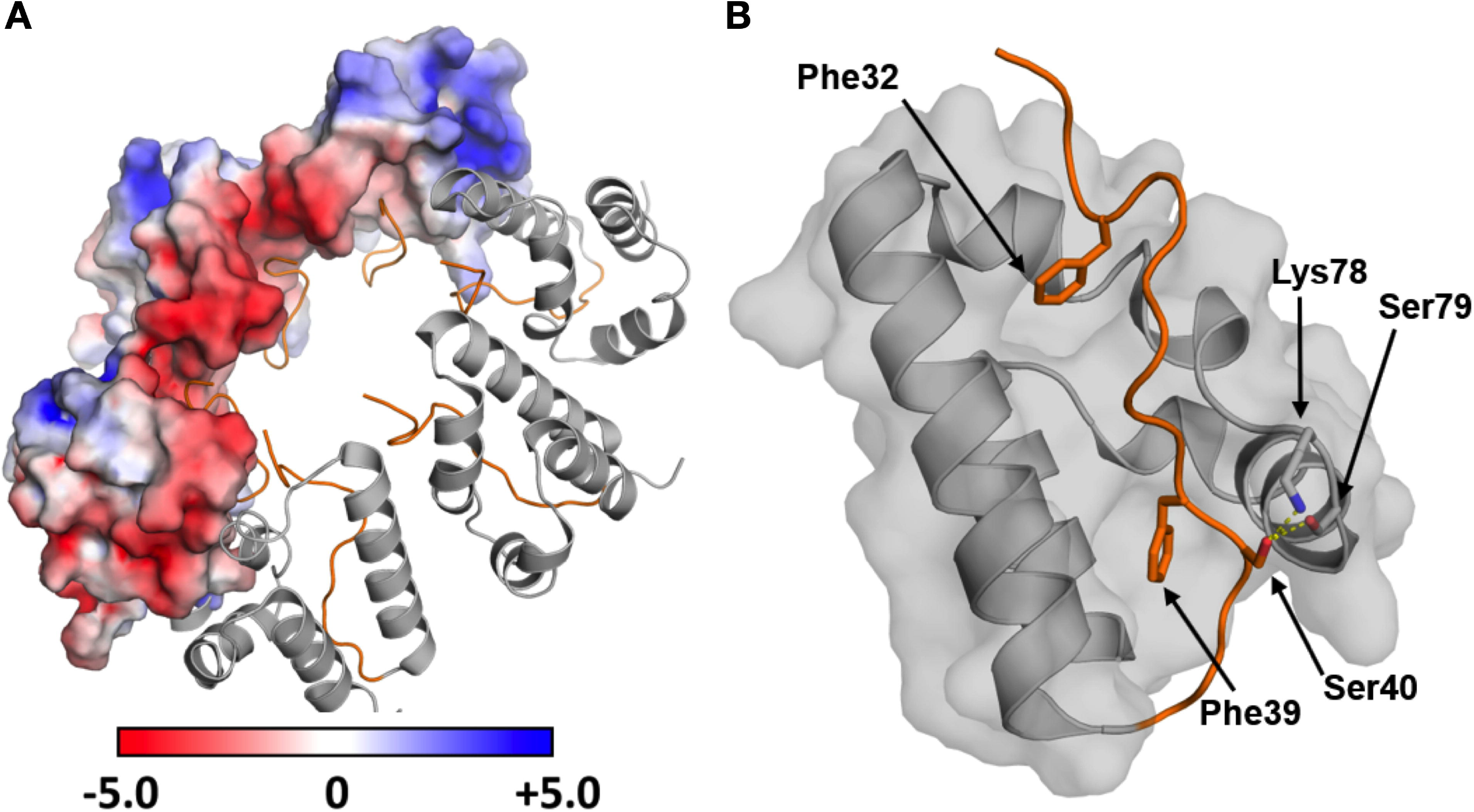
The N-terminal region preceding dArc-NL packs into the hydrophobic core of the domain and leads to the formation of the capsid hexamer. (A) The hexameric form of dArc2-NL observed in the capsid (PDB: 6TAQ; [20]), showing the electrostatic surface potential for half of the monomers. The splitting of α2 enables contact formation between the oppositely charged surfaces of each monomer. The N-terminal tail is showed in orange. (B) Residues contributing to the packing of the N-terminal tail (orange) into the capsid hexamer. Phe32 and Phe39 pack into two exposed pockets in the hydrophobic core. Further interactions are observed from Ser40 which hydrogen bonds directly with Lys78 and Ser79 in the α2 kink (interactions shown as yellow dashed lines).

In exploring the functional relevance of the domain swapped dArc2-NL dimer, we found that the overall fold of the dimer resembles retroviral proteins known to exhibit domain swapping. dArc2-NL resembles the flaviviral capsid C protein [56], which also forms domain-swapped dimers [57]. Despite low sequence similarity with the dengue virus 2 C protein (16.2%) and the core protein of the Kunjin subtype West-Nile virus (9.4%), the overall fold is surprisingly similar (Fig 4A-B) [56,58]. Both the dengue and West-Nile virus are enveloped RNA viruses, and the proteins are essential for the formation of the viral capsid. Despite different helix topology, the two proteins share fold similarity with dArc2-NL and have highly positive electrostatic surfaces, indicated by the authors to have a role in the binding of encapsulated genomic RNA [56,58]. Interestingly, in the crystal, the West-Nile virus core protein forms tetramers [58]. In the tetrameric form, long helices from each monomer (homologous to α2 in dArc2-NL) form a four-helix bundle subunit interface. This could be similar to the dArc2-NL tetrameric form in solution (see below). Moreover, the arrangement of dArc2-NL in the crystal bears some similarity to the domain-swapped dimer of the HIV CA lobe domain, induced by the deletion of a single residue [59].

**Fig 4.**
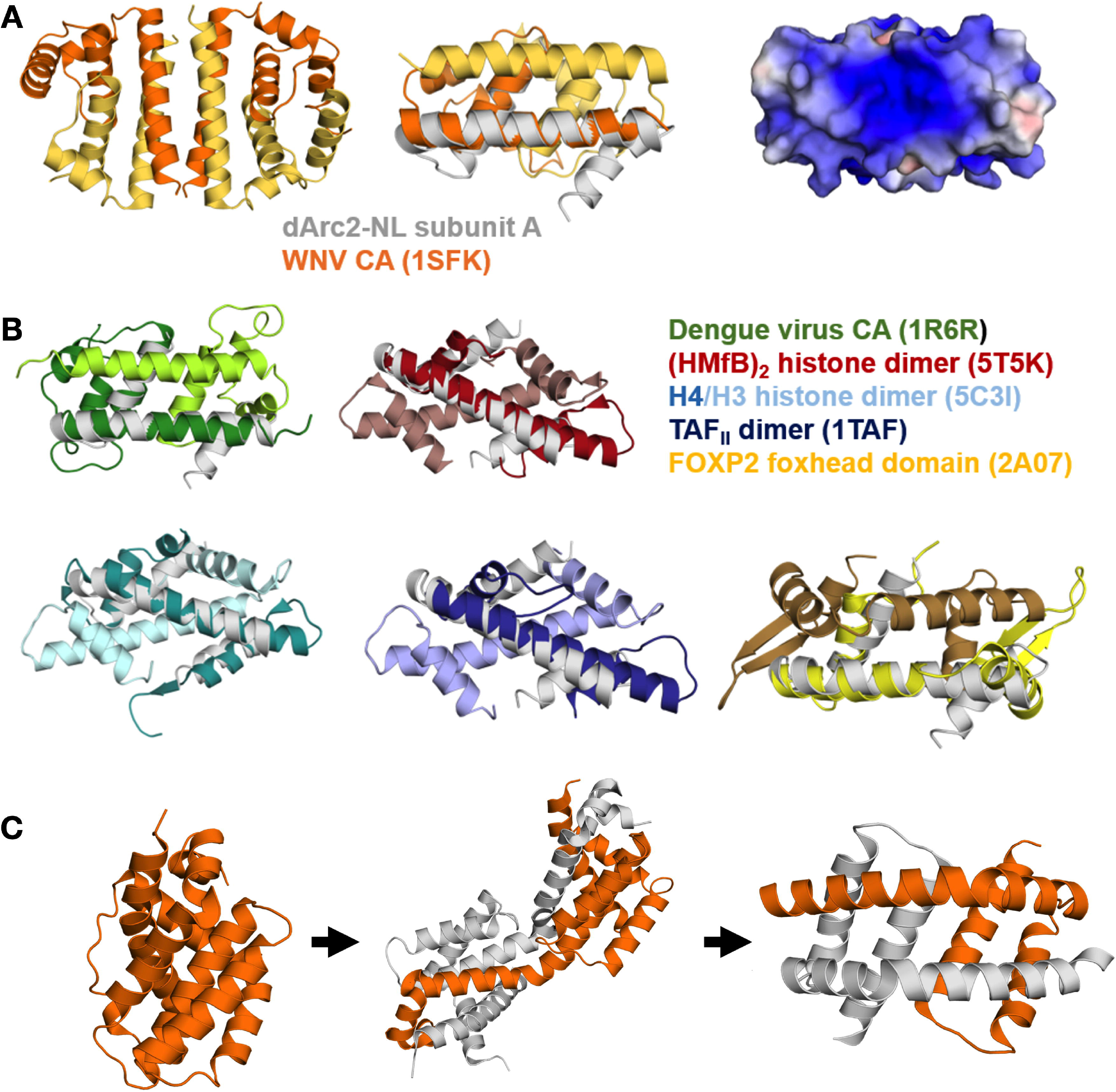
The dArc2-NL domain-swapped dimer resembles flaviviral coat proteins and DNA-binding proteins. (A) The tetrameric coat protein of the Kunjin subtype West-Nile virus (WNc), where the longest helix of each monomer (analogous to α2 of dArc2-NL) contributes to a four-helix bundle interface (PDB: 1SFK [58]) (left). Middle: a single dimer of the tetramer (yellow/orange) overlaid with subunit A from dArc2-NL (grey). Right: the electrostatic surface potential of a WNc dimer, which resembles that of dArc2-NL. (B) Structural comparison between the dArc2-NL and similar domain-swapping proteins. Shown are the retroviral Dengue virus CA (green; PDB: 1R6R [56]) and the DNA binding dimers of (HMfb)2 histone (red; PDB: 5T5K [60]), a dimer of histones H3 and H4 (cyan; PDB: 5C3I [61]), TAF_II_ transcription factor (blue; PDB: 1TAF [62]) and the foxhead domain of the FoxP2 transcription factor (yellow; PDB: 2A07 [63]). Each chain in a dimer is coloured with a different shade, and the dArc2-NL monomer is superimposed and shown in grey. (C) Domain swapping and conformational selection inthe apoptosis-induced BAK protein suggests a possible mechanism for dArc-2NL. Shown on the left is the inactive monomeric form of BAK (PDB: 2IMT [64]), which has an orthogonal bundle fold similar to Arc N-lobes. Binding of a BH3 domain causes partial unfolding and opening of the hinge region (middle, PDB: 4U2U [65]), which leads to the formation of the membrane-binding domain-swapped dimer (right, PDB: 4U2V [65]). Panel C is based on [66]. The two chains in the BAK dimer are coloured grey and orange.

Domain swapping of modular proteins emerges as a common theme in capsid-forming proteins. The dArc2-NL structure, with similarity to both retroviral and flaviviral capsid domain structures, shows that it is possible, via an extended helix, to transform the canonical capsid domain to a domain-swapped dimer. Whether such structures are related to the evolutionary history of capsid proteins, remains to be studied. It is interesting to note that the nucleocapsid protein from SARS coronavirus [67] also dimerises via domain swapping, while the sequence and structure are not similar to Arc.

In addition to viral capsid proteins, the structure of a single chain in the domain-swapped structure of dArc2-NL resembles the structure of a histone core protein monomer, as well as that of TATA box-binding protein-associated factors and the foxhead domain FoxP transcription factors (Fig 4B) [14,62,63,68,69]. The foxhead domain exist both as monomers and DNA binding domain-swapped dimers [63], which share significant fold topology with the dArc2-NL dimer. Moreover, upon replacement of a crucial alanine residue in the hinge region with proline (A39P), the foxhead domain lost all domain swapping ability [70]. The histone protein forms dimers, which combine to form tetramers and finally an octamer, to which DNA binds to form the nucleosome [71]. The histone and dArc2-NL dimer arrangements are different (Fig 4B), but the monomer structures are strikingly similar. This observation could be related to either the propensity of certain protein sequences to form domain-swapped structures or a functional similarity. The above observations on dArc2-NL are interesting in light of the histone mimicry by Dengue virus protein C [72], which interferes with host histones to inhibit nucleosome formation and gene transcription [9]. Whether such a mechanism could be important for Arc function, as mArc accumulates in the nucleus, associates with specific histone-modifying complexes, and is implicated in regulation of chromatin state and transcription [9,10,14,73] is a subject for future studies.

The same is true for the membrane binding core domains of BAK and BAX. BAK and BAX are members of the Bcl2 protein family and are important mediators of apoptosis. In its inactive form, BAK is monomeric and fully soluble, and the core domain has an orthogonal bundle fold. Upon activation, mediated by binding of certain BH3-only proteins into a hydrophobic groove in the core domain, the protein is partly unfolded, which leads to separation of the core and latch domains. The core domain then dimerizes to form amphipathic domain-swapped dimers [65,74]. These dimers can then further oligomerize and partition to the outer mitochondrial membrane where they bind and cause permeabilization, leading to the release of apoptosis factors such as cytochrome c into the cytosol [66]. Both the inactive and active forms of BAK show strikingly similar fold topology to the monomeric and dimeric state of the dArc2-NL, respectively. Additionally, the structure of the BAK intermediate, with the hinge region not fully open, might suggest a similar mechanism for conformational selection in dArc (Fig 4C). This might give valuable insight into the mechanism of domain swapping of dArc2-NL, in which interactions in the hydrophobic peptide binding groove seem of importance. Specific interactions in the groove might lead to dimerization, upon which the protein surface potential is rearranged to accommodate for nucleic acid or membrane binding.

### Evolutionary aspects of dArc2-NL domain swapping

Fig 5A shows the sequence entropy (opposite of conservation) of Arc NL and CL. In dArc2-NL, the most conserved residues are Ala81, Trp84, and Trp85, which sit on the hydrophobic side of the long α2 helix, corresponding to a conserved hydrophobic core in the domain family. Most interesting is Ser79 of dArc2, given its possible role in domain swapping. In all Arc-CL structures, the corresponding residue is a glycine in a β-turn, with a positive ⍰ angle. In the dArc2-NL structure, there is no β-turn, and the corresponding residue, Ser79, is in the middle of a long regular α-helix.

**Fig 5.**
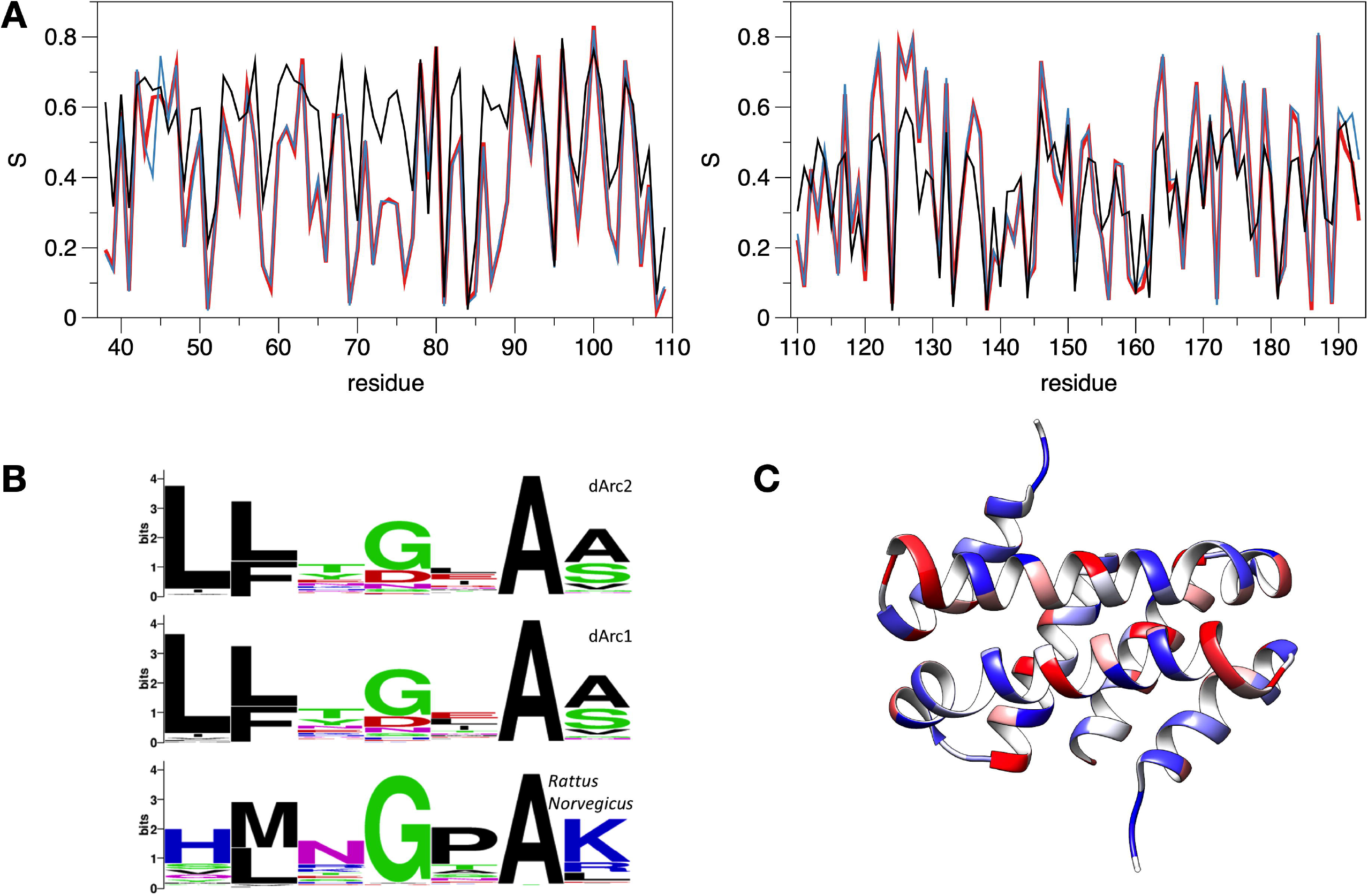
Sequence conservation analysis of the central lobe region. (A) Sequence variability in Arc N- and C-lobes (left and right, respectively). S is sequence entropy / variability. Numbering follows dArc2. Values are only shown for sites present in 50% or more of the sequences. dArc2 denotes 220 Arc2 homologues from Drosophila, dArc1+dArc2 a combined group of 250 homologues and dArc1+dArc2+Rattus has the group expanded to 699 sequences with homologues of the Rattus sequences. (B) Sequence logo for Ser79 of dArc2-NL compared to corresponding residues from dArc1 and Rattus norvegicus homologues. (C) Mapping of conservation onto the dArc2-NL dimer. Blue corresponds to conserved and red to non-conserved sites.

In evolutionary terms, Ser79 of dArc2-NL is an outlier, as clearly shown by a sequence logo of the region (Fig 5B). The window around Ser79 in dArc2 is `lfkSiav`, but whether one looks at homologues of dArc1, dArc2, or mArc, the site corresponding to Ser79 is most often a Gly, Asp, or Asn. These are common residues in a β turn [75], but Ser and Thr are also possible [76]; this changes the interpretation. In the dArc2-NL structure, one has a domain-swapped dimer and an α helix, where related structures have a β turn. Looking at the sequence homologues, the proteins have kept residues, which can adopt positive ⍰ angles and are likely to adopt turns. Looking at the conservation mapped onto the dArc2-NL structure, no clear cues are observed; rather, conserved residues are evenly dispersed along the folded structure (Fig 5C). In the structures of the dArc1 and dArc2 capsid [20] as well as the crystal structure of a longer dArc1 construct [55], the NL has the canonical fold without domain swapping. These features imply that the domain-swapped dArc2-NL structure might be due to crystallization of one domain alone, but it confirms the general capability of CA domains to dimerize through different modes, including domain swapping [53,54,59,77].

### Structures of dArc C-lobes

Both dArc1 and dArc2 CL crystallized as homodimers (Fig 6A-B, S2, S3), and each monomer consists of five helices in an orthogonal bundle fold. The structures are highly similar (Fig 6C). The dimer interface is in both cases formed by α1 and α3 from each monomer, and the total buried surface area at the interface is similar, ~1400 Å^2^ (Fig 6D). Both interfaces contain four hydrogen bonds and four salt bridges. The interface is conserved, displaying only three conservative replacements (A125/L170/F172 in dArc1 to S112/F157/Y159 in dArc2). The dArc-CL dimers resemble the domain in the dArc capsids (Fig 6E), with an all-atom RMSD of 1.75 Å for dArc1 and 1.13 Å for dArc2. The intact CA domain of dArc1 is dimeric in solution [55]. Conservation of the dimer interface suggests a vital function of this mode of oligomerization in dArc function, both as a capsid and dimeric in solution.

**Fig 6.**
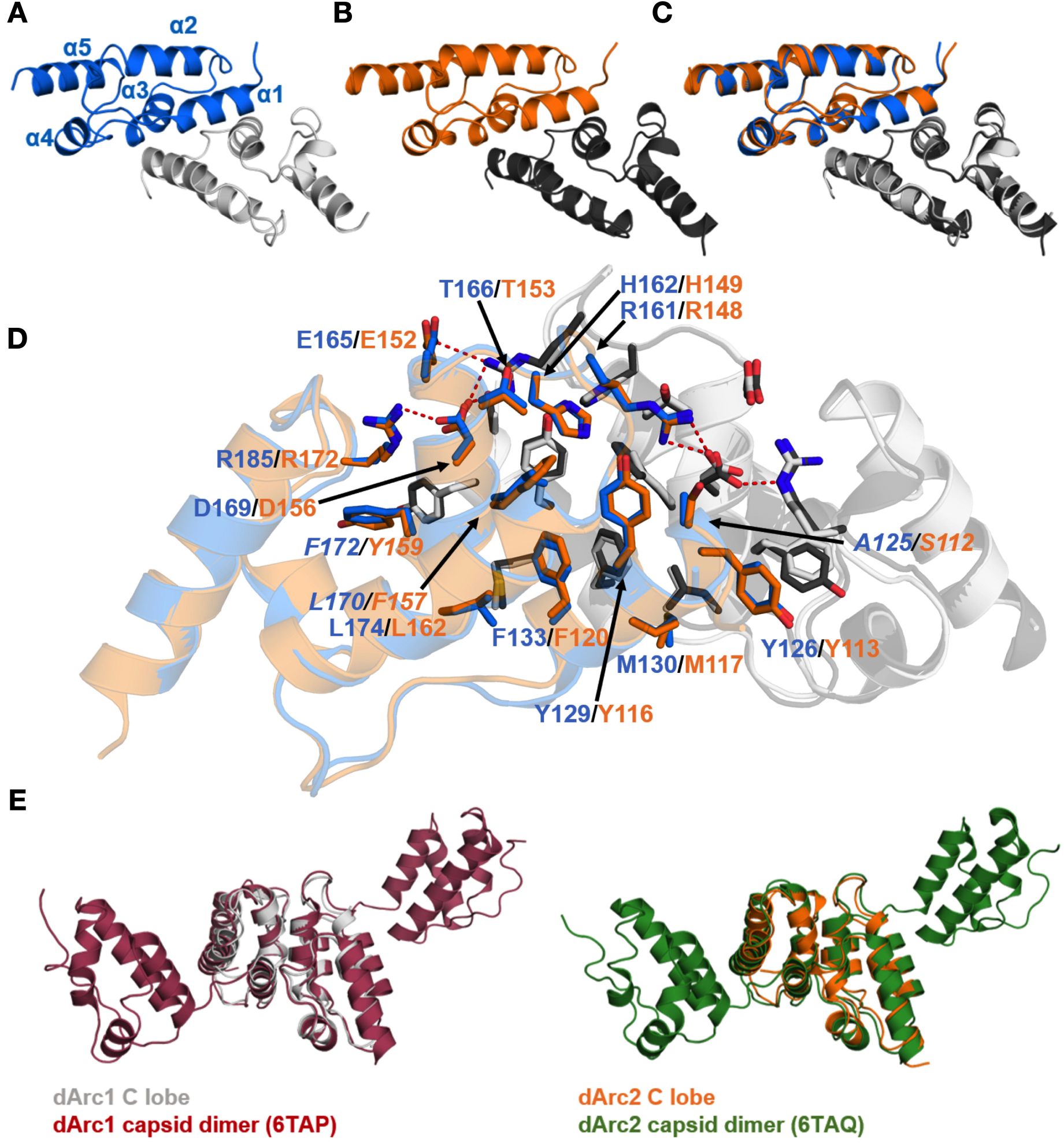
Crystal structures of the dArc1 and dArc2 C-lobes reveal dimeric orthogonal bundles. (A) dArc1-CL. (B) dArc2-CL. (C) The two structures, which deviate with an all-atom RMSD of 0.48 Å, aligned. (D) Residues contributing to the dimer interface in dArc2-CL. dArc1-CL residues are marked in blue, and residues of dArc2-CL are marked in orange. Altering residues are indicated in italics. All residues contributing to the dimer interface are highly conserved, with the exception of A125 (dArc1) which corresponds to S112 (dArc2). Polar interactions are shown with red dashed lines. (D) A comparison of the dArc C-lobe crystal structures with the same domains in the viral-like capsids formed by the protein. Both the dArc1 and dArc2 C-lobes closely resemble their counterparts in the capsids, with an all-atom RMSD of 1.75 Å^2^ and 1.13 Å^2^, respectively.

The dArc CL domains are dimeric also in solution (see below). This behaviour of the dArc C-lobes is different to the monomeric mArc C-lobe [17], while both share the same core structure [12]. The dimer interface in dArc-CL, which corresponds to that in retroviral CA-CTD [54], contains mainly hydrophobic interactions; half of these hydrophobic residues are polar in the rat Arc CL, and the first helix of the dArc-CL, a major part of the dimer interface, is tilted away in mArc, explaining the monomeric state of mArc-CL in solution [17].

The crystal structures of the dArc C-lobes resemble those of mArc and retroviral capsid proteins (Fig 7A). The dimerization of the retroviral CA-CTD is similar to that of dArc-CL; α1 and α3 of each five-helix bundle contribute to the subunit interface (Fig 7B). However, in both HIV and bovine leukemia virus (BLV) CTDs, the N-terminal segment differs from Arc, consisting of a seven- and six-helix orthogonal bundle, respectively. HIV forms elongated conical capsids, and HIV-1 CA assembles spontaneously into helical tubes in vitro [78]. Thus, despite high similarity of individual domains within the fold family, the assembly mechanisms into larger structures may be different and depend on additional domain modules in the corresponding protein.

**Fig 7.**
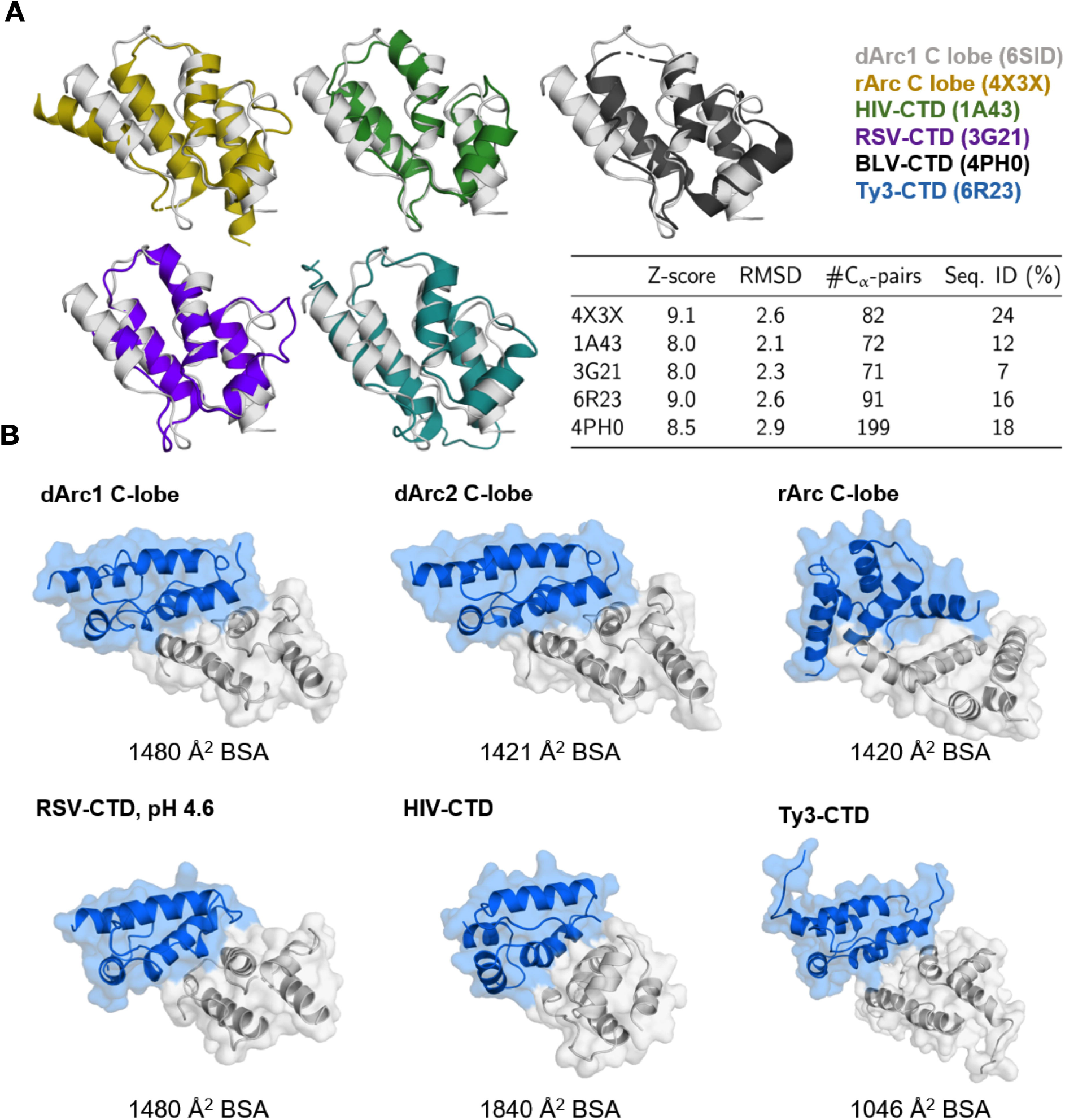
The structure of dArc-CL resembles that of mArc and retroviral capsid proteins. (A) Structures similar to dArc1-CL. dArc1-CL (grey) is shown aligned with crystal structures of the rat Arc C-lobe (yellow; PDB: 4X3X) [12], HIV C-terminal domain (green; PDB:1A43) [79], bovine leukemia virus (BLV) C-terminal domain (black; PDB:4PH0) [80], the rous sarcoma virus (RSV) C-terminal domain crystallized at pH 4.6 (purple; PDB: 3G21) [81], and the C-terminal domain of the Ty3 retrotransposon capsid (cyan; PDB: 6R23) [82]. Also shown are the scoring criteria obtained from the Dali server. (B) Comparison of CT dimerization. Shown are the dimer interfaces of the structural homologues in (A) and dArc2-CL, as calculated by PISA [37] from the crystalline states, apart from the BSV-CTD, which was not dimeric. Buried surface area (BSA) of each interface is shown below each structure.

Both the intact mArc-CT and the mArc-CL alone are monomeric in solution [12,17,83]. The crystal structure of the rat Arc CL suggests a monomeric state [12], and a dimer similar to dArc-CL cannot be found in the crystal symmetry. However, a likely dimeric state of the protein was found in the crystal lattice by PISA (Fig 7B). Despite the high structural similarity to both the dArc C-lobes, oligomerization differs in mArc. In this putative dimer, the interface is formed by α1 and α2 of each monomer. The total buried surface area at the interface is similar to both dArc1-CL and dArc2-CL, being composed of 75 van der Waals and π-π contacts, 2 hydrogen bonds, and 4 salt bridges.

Conservation within the Arc-CL (Fig 5A) raises some questions. The most conserved residue (Gln124 in dArc2) is structurally important, forming hydrogen bonds and contacts with many neighbours, including the conserved residues Phe133 and Met162. Arg138 and Asp151 are surface-exposed, but highly conserved in both insects and mammals. Therefore, they could be central in a network of salt bridge interactions on the CL surface. It is likely that such conserved residues are required for the correct folding of the Arc lobe structure.

### All dArc lobe domains are oligomeric in solution

The structure and oligomeric state of the dArc lobe domains were analyzed in solution by SEC-MALS, SAXS, and CD (Fig 8, Table 2). Both dArc1-CL and dArc2-CL are compact and slightly elongated, fitting the crystallographic dimers (Fig 8A-B). dArc1-NL is similar, being the size of a dimer. dArc2-NL is twice the size of dArc1-NL in solution, suggesting a tetramer. The details of the latter arrangement are currently unknown, since no symmetric tetrameric assemblies can be deduced from the crystal structure, but the assembly could be similar to the West Nile virus C protein [58]. The tetrameric C protein resembles the dArc2-NL (Fig 4A), in that it is a dimer of domain-swapped dimers similar to the dArc2-NL homodimer. SAXS data for dArc2-NL in solution fit the structure of a similar tetrameric assembly (Fig 8C-D). Hence, it is possible that the observed dArc2-NL tetramers in solution assemble in the same way as those for the C protein.

**Table 2.**
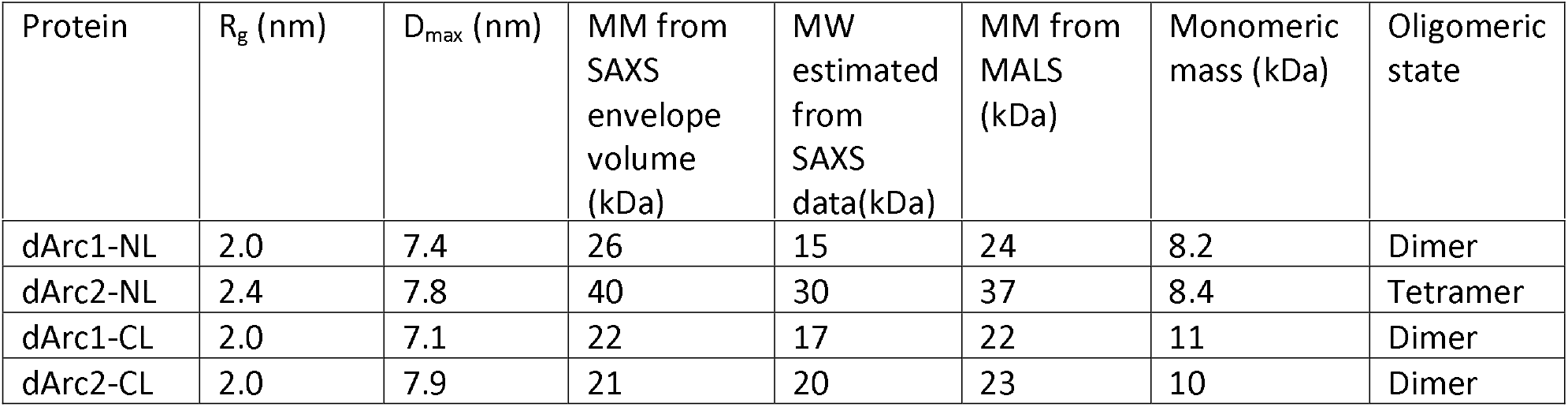
Dimensions and oligomeric state for different dArc constructs.

**Fig 8.**
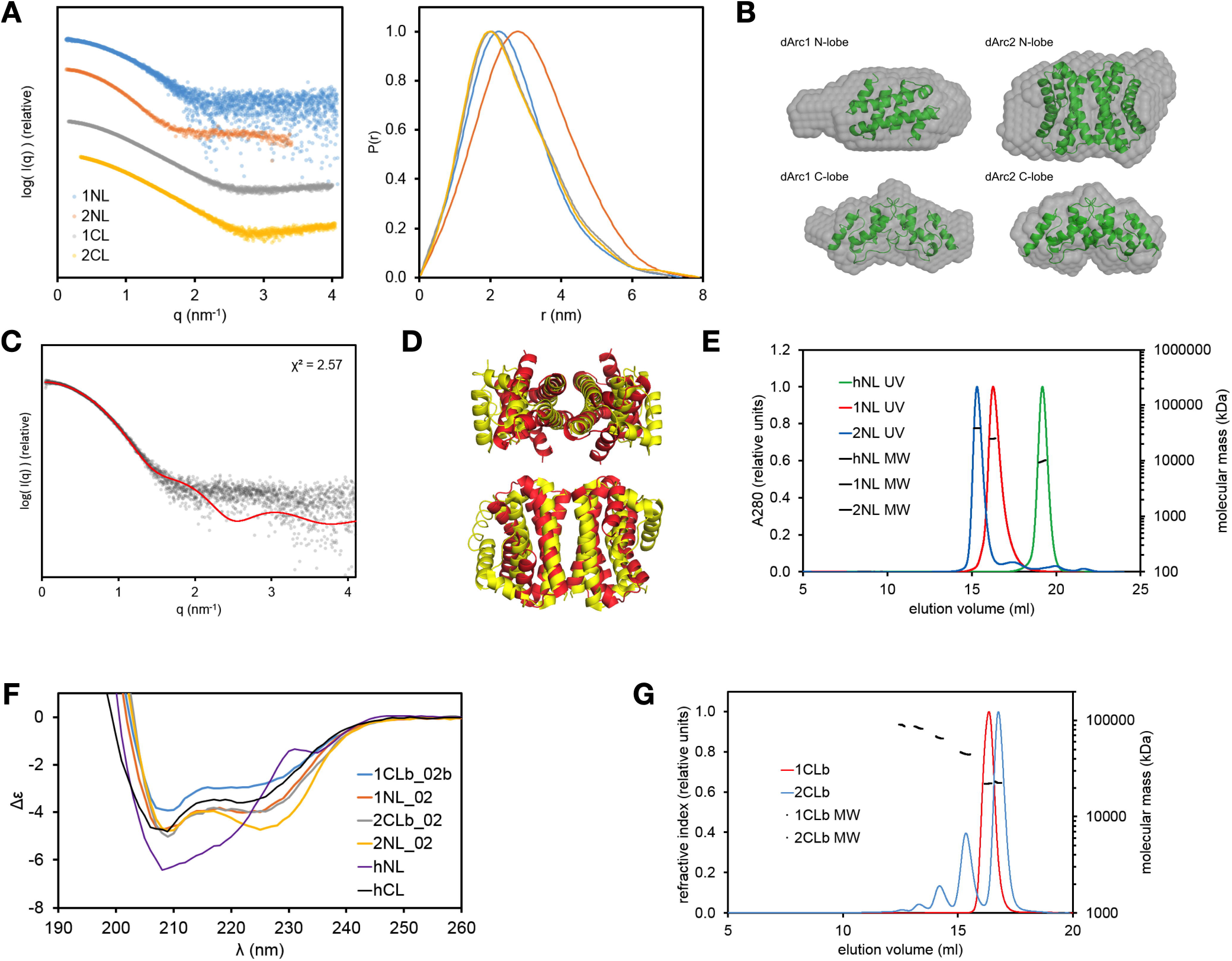
Solution structures of dArc lobe domains. (A) SAXS scattering plots of dArc lobes in solution (left) and distance distribution plots (right). (B) Dammin models of dArc lobes with corresponding structures located inside. The dimeric structures of the C-lobes inside the C-lobe Dammin models and the assumed tetrameric structure of dArc2 N-lobe in the dArc2 N-lobe Dammin model, and the structure of dimeric dArc2 N-lobe in the Dammin model of dArc1 N-lobe. (C) SAXS data for of dArc2-NL (dots) with crysol fit using the possible tetrameric structure of dArc2 N-lobe based on the West Nile virus tetramer structure (red) seen in panel (D). (D) Tetrameric organisation of the West Nile virus protein C (yellow) and aligned structure of two dArc2 N-lobe dimers (red) showing a possible tetrameric structure. (E) SEC-MALS on human and Drosophila N-lobes. (F) CD data of dArc lobes and human Arc lobes. (G) SEC-MALS on human and Drosophila C-lobes.

Arc-NL domains have various oligomeric states. mArc-NL is monomeric in solution [17], whereas the dArc-NL forms dimers (dArc1) and tetramers (dArc2) (Fig 8E). While the crystal structure of dArc2-NL shows a dimer, a tetrameric assembly is not present in the crystal. This is remarkable, as the sequences of dArc1-NL and dArc2-NL are very similar (Fig 9A). The dArc2-NL dimer surface is electrostatically polarised (Fig 2). By threading the sequence of dArc1 onto the dArc2-NL structure, the varying residues are mainly located on the surface of the long helices (α2 and α2’, Fig 2), suggesting that this region is responsible for dArc2-NL tetramerization. Presumably, a tetramer of dArc2-NL is achieved either by formation of a four-helix interface bundle, similar to the West-Nile virus coat protein (Fig 4A) or via interaction of the contrasting surface potentials on each side of the dimer. The lack of this electrostatic polarisation in dArc1-NL could be linked to oligomerisation (Fig 9B). Furthermore, additional polar interactions are observed at the interface of a dArc1-NL domain-swapped homology model (Fig 9C), compared to the dArc2-NL interface, which only consists of nonpolar contacts. These different interactions could also be linked to dArc-NL isoform-specific oligomerisation.

**Fig 9.**
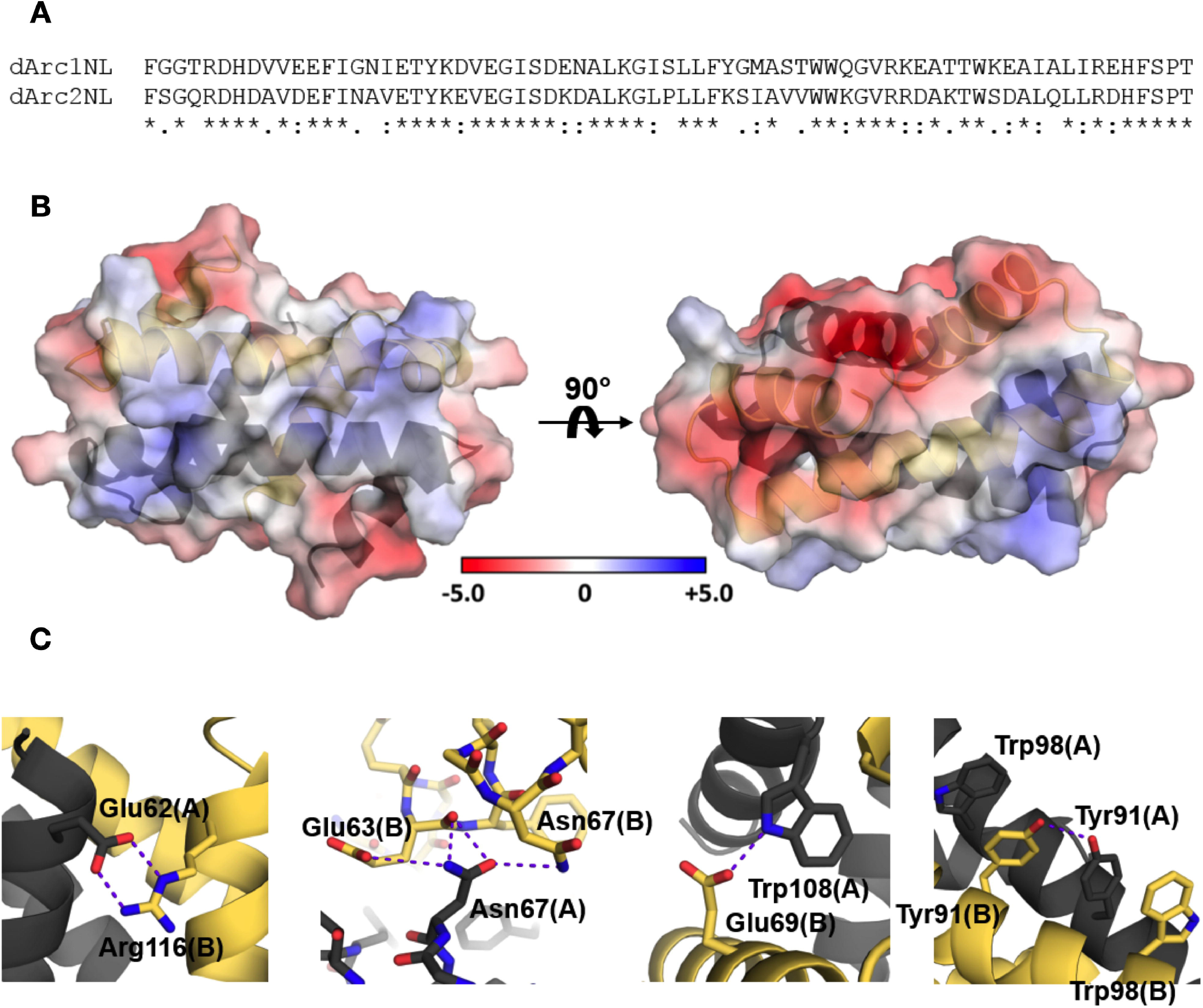
dArc1-NL homology model. (A) Sequence alignment between dArc1-NL and dArc2-NL. (B) The homology model of dArc1-NL displays contrasting electrostatic surface potential, where the highly positive character of dArc2-NL along α2 and α2’ (Fig 2) is replaced with a more modest surface potential. (C) Additional monomer-monomer interactions observed in the dArc1-NL model, not observed in the dArc2-NL crystal structure. Polar interactions are shown with purple dashed lines.

CD spectroscopy showed that all four dArc lobes are α-helical (Fig 8F), with some variations in spectral shape and amplitude. dArc2-NL has a higher 222-to-208-nm ratio compared to dArc1-NL. This could be related to differences in dimerization (domain swapping) or tetramer formation. Tetramerization may involve interactions between the long helices of dArc2-NL, and coiled-coil interactions increase the 222-to-208-nm ratio [84–86]. CD spectra of the dArc C-lobes are similar but differ in intensity, suggesting that dArc1-CL is less folded in solution, despite the very similar crystal structures. In line with the CD data, SEC showed (Fig 8G) a higher hydrodynamic radius for dArc1-CL. Kratky plots also indicate that dArc1-CL is more flexible than dArc2-CL. The CD spectrum of monomeric hArc-CL is similar to dArc-CL but shows less helical structure (Fig 8F). The monomeric hArc-NL has unique CD features, possibly arising from interactions between aromatic side chains. Hence, different methods for following protein folding and shape suggest that each of the dArc lobes has unique properties compared to each other and to homologues.

### Understanding higher-order oligomerization

We determined the crystal structure of three of the four dArc lobe domains: both C-lobes and dArc2-NL; a crystal structure for dArc1-NL could not be obtained. However, the high sequence similarity between dArc1-NL and dArc2-NL (Fig 9A) suggests that the structure of dArc1-NL is similar to dArc2-NL. Furthermore, secondary structure analysis using CD shows similar spectra for both proteins (Fig 8F), and the structure of the dArc2-NL dimer fits well with the SAXS data for dArc1-NL (Fig 8C-D). However, other dimeric arrangements for dArc1-NL are possible. In this respect, it is interesting to note the loss of a conserved Gly residue in the canonical ⍰2-⍰3 loop in dArc2-NL, which could be related to the extension of the dArc2-NL helix. Replacement of a Gly residue in such a loop is a common means to induce domain swapping [87,88]. The recent crystal structure of dArc1-CT containing both lobes showed dArc1-NL in the orthogonal bundle fold, being similar to mArc-NL [55]. No significant interactions were observed between the NL and CL in dArc1-CA, and interlobal interactions are an unlikely cause of the different fold. Neither CD nor SAXS can determine if dArc1-NL has the same domain-swapped structure as dArc2-NL or a non-domain-swapped dimer as seen for dArc-CL.

All four lobe domains of dArc are homo-oligomeric in solution. In full-length dArc, the NL is connected with the CL, and to test for interactions between the dArc N- and C-lobes, we mixed the individual lobes and looked for complexes using SEC. No new complexes were observed (data not shown), indicating that the isolated NL and CL do not interact with high affinity. However, when both lobes are within the same polypeptide chain, larger assemblies do form – reflected by the insolubility of the corresponding constructs and the ability of dArc to form capsids [15,20].

### dArc sequence properties

Arc may not be a universal protein, but it is truly ancient. Related proteins appear in eukaryotes, from insects to fungi and plants [89]. At the same time, Arc-like proteins are coded for by the Ty3/gypsy transposons, and its relatives appear in viral capsids. This means the domain is widespread because of duplications and movements within and between genomes, rather than its age. This invites some speculation about the history of Arc, or at least the history of the N- and C-lobes.

The NL and CL are sequence-related, suggesting a duplication. They are related to viral capsid (Gag) proteins, but the Gag protein in flavi- and other viruses has only one unit, or lobe. One might expect to see either the NL or CL by itself in some cellular organism. A long, iterated search starting from either *Drosophila* or *Rattus* full sequences only gives proteins with both NL and CL, even amongst distantly related proteins from plants. This is not surprising. Database scores are such that a long weak similarity will score higher than a short hit, and one will see proteins with both lobes. The correct procedure is to do a comprehensive database search starting from the CL, retrieve and align full length sequences, and see if any are missing the NL. This should then be repeated starting from the NL. Unfortunately, this does not give a clear result. Starting from the rat Arc-CL, one can collect a set of 706 sequences with an *e*-value ≤ 1.4×10^−5^. The set runs from mammals to insects and even the first homologues from plants (*Oryza sativa* and *Nicotiana tabacum*). We find 26 sequences with an incomplete NL. More than a third (9) of these are annotated as partial sequences. Of the remaining sequences, none are confirmed to exist, and there is no clear domain boundary in the alignment. Similar results are obtained starting from an NL sequence. This does, however, not prove conclusively that the NL and CL are always found together in eukaryotes, and we might even expect some related transposon sequences to be lurking with only one lobe.

In eukaryotes from mammals to insects and plants, the overwhelming majority of Arc-like proteins have both NL and CL. This has implications for the Arc evolutionary history. It appears that a functional protein has both lobes, and there are no clear examples of a system with two copies of just an NL or CL. The obvious interpretation is that after a duplication event, the two halves adopted different roles and sites in the two halves have experienced differential evolutionary pressure, as suggested by the conservation plots (Fig 5A) and the observation [20] that both dArc lobes are necessary for capsid assembly.

It was suggested that Arc entered the realm of animals *via* two separate events [16]. The earlier work relied on a small set of DNA sequences. When we align larger sets of proteins including mammals, insects, birds, and plants, one always finds the insect sequences forming a close group including both dArc1 and dArc2. Even a cursory look at the alignment (Fig 9A) shows the similarity of dArc1 and dArc2. This leads to a parsimonious interpretation. The combined NL and CL protein was inserted once, leading to the tandem arrangement nearly always seen. This protein has had much time to evolve between insects and mammals. At a much later stage, a duplication led to the multiple copies seen in insects.

### Functional considerations

Arc is critical to the nervous system, but the protein fold is related to capsid proteins and, at least at the sequence level, even to proteins found in plants. A functional property of the mArc-NL is a peptide binding site, shown to interact with several proteins [12]. Protein ligand binding could be a way of regulating Arc oligomerization [83], its function in the PSD organization, and/or capsid assembly. Using ITC, we tested if the dArc N-lobes bind the stargazin peptide like the mArc-NL (Fig 10) [12]. The peptide binds to hArc-NL, but not to the two dArc N-lobes, and the peptide binding site is not conserved. In the domain-swapped structure of dArc2-NL, the putative peptide binding site would be buried within the fold. Whether dArc N-lobes bind other peptides/proteins, and how this process might affect capsid formation, remains to be studied. Given that the dArc N-lobe forms the pentamers and hexamers in the capsid [20], and that the site corresponding to the mArc peptide binding site is in the middle of these assemblies, it seems likely that protein ligand binding to the same site would be mutually exclusive with capsid formation. Such aspects remain to be studied both for dArc and mArc.

**Figure 9.**
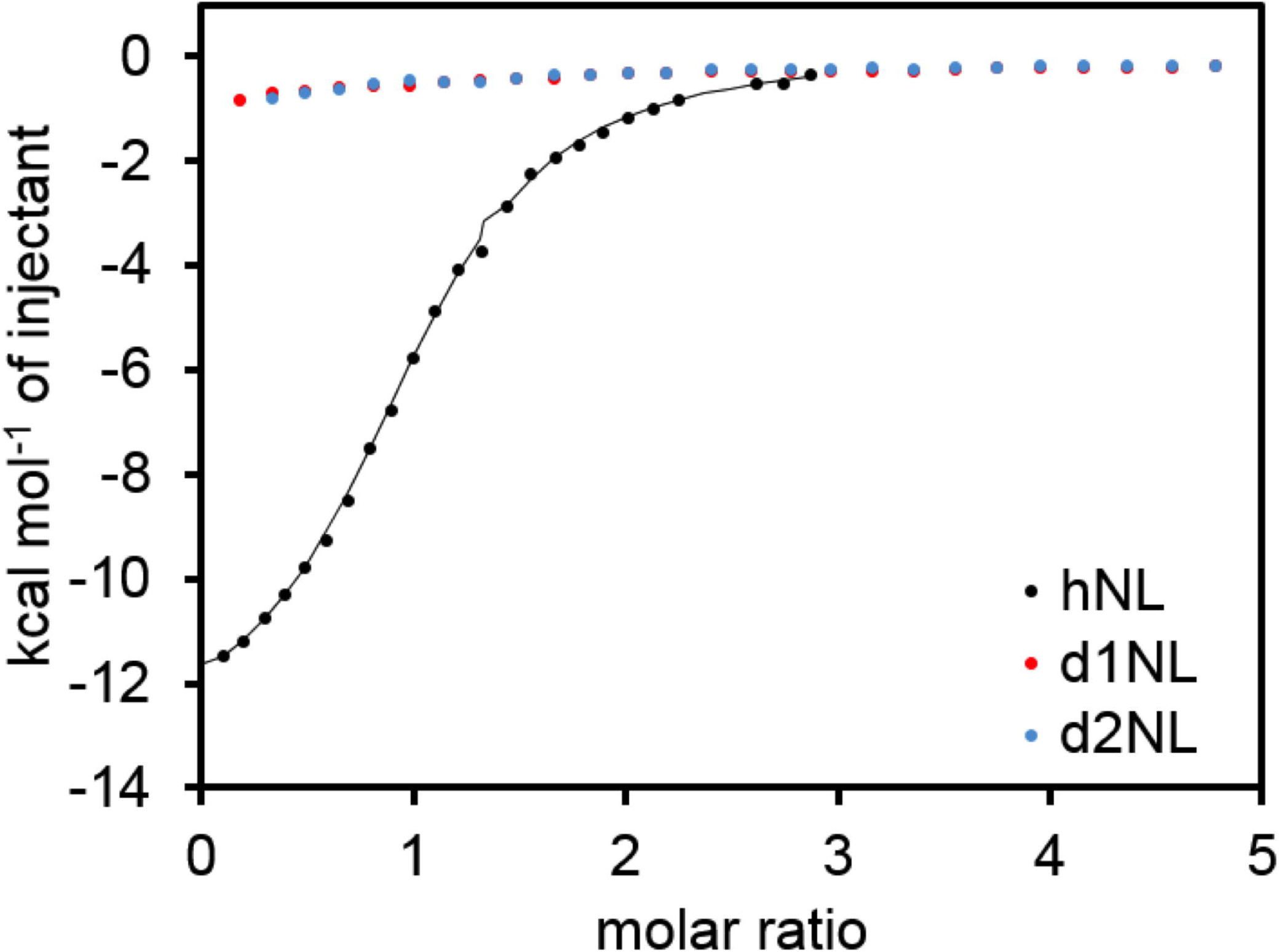
Binding of human and Drosophila Arc N-lobes to a Stargazin peptide.

## Conclusions

We have shown that both lobes of dArc1 and dArc2 are oligomeric in solution, the mechanism of which was further explored in atomic detail. The isolated lobes of the C-terminal domain of mammalian Arc do not exhibit the same propensity for oligomerization [17]. Absent in both dArc isoforms is the N-terminal domain, which is involved in capsid formation and likely mediates interactions of mammalian Arc with negatively charged membranes [17,18]. Therefore, oligomerization of the dArc lobes likely represents functional compensation for the lack of the N-terminal domain. In accordance with this, dimerization of the C-lobe observed here is identical to that observed in a recent crystal structure of dimeric full-length dArc1 [55] and structures of the capsids formed by dArc [20]. Our C-lobe structures showed striking homology with retroviral CA proteins, further supporting theories on the retroviral origin of Arc. However, the homologous retroviruses form capsids of greatly varying morphology. This suggests a decisive role of the N-terminal lobe, and its mechanism of oligomerization, in the formation of these viral capsids.

Furthermore, we presented a novel dimeric state of dArc2-NL. This domain-swapped dimer may have a role in the non-capsid functions of dArc. It shares structural similarity with nucleotide- and membrane-interacting proteins, suggesting a similar function for the fold. How homotetramerization of this lobe, which we observed in solution, might affect its function, remains to be studied. Overall, the strikingly different behaviour of the purified lobe domains from dArc and mArc points towards different mechanisms in their molecular function and oligomeric assembly.

## Acknowledgements

Parts of this research were carried out on beamline P11 at DESY, a member of the Helmholtz Association (HGF). We wish to thank EMBL/DESY for access to beamline P12 and acknowledge Diamond Light Source for time on Beamlines I03 and B21 under Proposal MX18666. We would like to thank all beamline staff for assistance during the experiments. The research leading to this result has been supported by the project CALIPSOplus under the Grant Agreement 730872, as well as by iNEXT, grant number 653706, from the EU Framework Programme for Research and Innovation HORIZON 2020. This work was supported by a Research Council of Norway TOPPFORSK grant (249951) to CRB and a grant from the Meltzer Foundation to PK. The authors wish to express their gratitude to Ju Xu for technical assistance.

## Supporting information

S1 Table. The expression constructs used in the current study.

S1 Fig. Crystal packing in dArc2-NL.

S2 Fig. Crystal packing in dArc1-CL.

S3 Fig. Crystal packing in dArc2-CL.

## Notes

### Competing Interest Statement

The authors have declared no competing interest.

## References

1. Bramham CR, Alme MN, Bittins M, Kuipers SD, Nair RR, Pai B et al. (2010) The Arc of synaptic memory. Exp Brain Res 200: 125–140.

2. Shepherd JD, Bear MF (2011) New views of Arc, a master regulator of synaptic plasticity. Nat Neurosci 14: 279–284.

3. Chowdhury S, Shepherd JD, Okuno H, Lyford G, Petralia RS, Plath N et al. (2006) Arc/Arg3.1 interacts with the endocytic machinery to regulate AMPA receptor trafficking. Neuron 52: 445–459.

4. Rial Verde EM, Lee-Osbourne J, Worley PF, Malinow R, Cline HT (2006) Increased expression of the immediate-early gene arc/arg3.1 reduces AMPA receptor-mediated synaptic transmission. Neuron 52: 461–474.

5. Shepherd JD, Rumbaugh G, Wu J, Chowdhury S, Plath N, Kuhl D et al. (2006) Arc/Arg3.1 mediates homeostatic synaptic scaling of AMPA receptors. Neuron 52: 475–484.

6. Messaoudi E, Kanhema T, Soulé J, Tiron A, Dagyte G, da Silva B et al. (2007) Sustained Arc/Arg3.1 synthesis controls long-term potentiation consolidation through regulation of local actin polymerization in the dentate gyrus in vivo. J Neurosci 27: 10445–10455.

7. Peebles CL, Yoo J, Thwin MT, Palop JJ, Noebels JL, Finkbeiner S (2010) Arc regulates spine morphology and maintains network stability in vivo. Proc Natl Acad Sci U S A 107: 18173–18178.

8. Zhang H, Bramham CR (2020) Arc/Arg3.1 function in long-term synaptic plasticity: Emerging mechanisms and unresolved issues. Eur J Neurosci

9. Korb E, Wilkinson CL, Delgado RN, Lovero KL, Finkbeiner S (2013) Arc in the nucleus regulates PML-dependent GluA1 transcription and homeostatic plasticity. Nat Neurosci 16: 874–883.

10. Wee CL, Teo S, Oey NE, Wright GD, VanDongen HM, VanDongen AM (2014) Nuclear Arc Interacts with the Histone Acetyltransferase Tip60 to Modify H4K12 Acetylation(1,2,3). eNeuro 1:

11. Jackson AC, Nicoll RA (2011) Stargazing from a new vantage--TARP modulation of AMPA receptor pharmacology. J Physiol 589: 5909–5910.

12. Zhang W, Wu J, Ward MD, Yang S, Chuang YA, Xiao M et al. (2015) Structural basis of arc binding to synaptic proteins: implications for cognitive disease. Neuron 86: 490–500.

13. Zhao Y, Chen S, Yoshioka C, Baconguis I, Gouaux E (2016) Architecture of fully occupied GluA2 AMPA receptor-TARP complex elucidated by cryo-EM. Nature 536: 108–111.

14. Nikolaienko O, Patil S, Eriksen MS, Bramham CR (2018) Arc protein: a flexible hub for synaptic plasticity and cognition. Semin Cell Dev Biol 77: 33–42.

15. Ashley J, Cordy B, Lucia D, Fradkin LG, Budnik V, Thomson T (2018) Retrovirus-like Gag Protein Arc1 Binds RNA and Traffics across Synaptic Boutons. Cell 172: 262–274.e11.

16. Pastuzyn ED, Day CE, Kearns RB, Kyrke-Smith M, Taibi AV, McCormick J et al. (2018) The Neuronal Gene Arc Encodes a Repurposed Retrotransposon Gag Protein that Mediates Intercellular RNA Transfer. Cell 173: 275.

17. Hallin EI, Eriksen MS, Baryshnikov S, Nikolaienko O, Grødem S, Hosokawa T et al. (2018) Structure of monomeric full-length ARC sheds light on molecular flexibility, protein interactions, and functional modalities. J Neurochem 147: 323–343.

18. Eriksen MS, Nikolaienko O, Hallin EI, Grødem S, Bustad HJ, Flydal MI et al. (2020) Arc self-association and formation of virus-like capsids are mediated by an N-terminal helical coil motif. FEBS J

19. Zhang W, Chuang YA, Na Y, Ye Z, Yang L, Lin R et al. (2019) Arc Oligomerization Is Regulated by CaMKII Phosphorylation of the GAG Domain: An Essential Mechanism for Plasticity and Memory Formation. Mol Cell 75: 13–25.e5.

20. Erlendsson S, Morado DR, Cullen HB, Feschotte C, Shepherd JD, Briggs JAG (2020) Structures of virus-like capsids formed by the Drosophila neuronal Arc proteins. Nat Neurosci

21. van den Berg S, Löfdahl PA, Härd T, Berglund H (2006) Improved solubility of TEV protease by directed evolution. J Biotechnol 121: 291–298.

22. Raasakka A, Myllykoski M, Laulumaa S, Lehtimäki M, Härtlein M, Moulin M et al. (2015) Determinants of ligand binding and catalytic activity in the myelin enzyme 2’,3’-cyclic nucleotide 3’-phosphodiesterase. Sci Rep 5: 16520.

23. Burkhardt A, Pakendorf T, Reime B, Meyer J, Fischer P, Stübe N et al. (2016) Status of the crystallography beamlines at PETRA III. The European Physical Journal Plus 131: 56.

24. Kabsch W (2010) XDS. Acta Crystallogr D Biol Crystallogr 66: 125–132.

25. Bibby J, Keegan RM, Mayans O, Winn MD, Rigden DJ (2012) AMPLE: a cluster-and-truncate approach to solve the crystal structures of small proteins using rapidly computed ab initio models. Acta Crystallogr D Biol Crystallogr 68: 1622–1631.

26. Xu D, Zhang Y (2012) Ab initio protein structure assembly using continuous structure fragments and optimized knowledge-based force field. Proteins 80: 1715–1735.

27. Krissinel E, Uski V, Lebedev A, Winn M, Ballard C (2018) Distributed computing for macromolecular crystallography. Acta Crystallogr D Struct Biol 74: 143–151.

28. Potterton L, Agirre J, Ballard C, Cowtan K, Dodson E, Evans PR et al. (2018) CCP4i2: the new graphical user interface to the CCP4 program suite. Acta Crystallogr D Struct Biol 74: 68–84.

29. Panjikar S, Parthasarathy V, Lamzin VS, Weiss MS, Tucker PA (2005) Auto-rickshaw: an automated crystal structure determination platform as an efficient tool for the validation of an X-ray diffraction experiment. Acta Crystallogr D Biol Crystallogr 61: 449–457.

30. Sheldrick GM (2008) A short history of SHELX. Acta Crystallogr A 64: 112–122.

31. McCoy AJ, Grosse-Kunstleve RW, Adams PD, Winn MD, Storoni LC, Read RJ (2007) Phaser crystallographic software. J Appl Crystallogr 40: 658–674.

32. Cowtan K (2010) Recent developments in classical density modification. Acta Crystallogr D Biol Crystallogr 66: 470–478.

33. Cowtan K (2012) Completion of autobuilt protein models using a database of protein fragments. Acta Crystallogr D Biol Crystallogr 68: 328–335.

34. Afonine PV, Grosse-Kunstleve RW, Echols N, Headd JJ, Moriarty NW, Mustyakimov M et al. (2012) Towards automated crystallographic structure refinement with phenix.refine. Acta Crystallogr D Biol Crystallogr 68: 352–367.

35. Emsley P, Lohkamp B, Scott WG, Cowtan K (2010) Features and development of Coot. Acta Crystallogr D Biol Crystallogr 66: 486–501.

36. Williams CJ, Headd JJ, Moriarty NW, Prisant MG, Videau LL, Deis LN et al. (2018) MolProbity: More and better reference data for improved all-atom structure validation. Protein Sci 27: 293–315.

37. Krissinel E, Henrick K (2007) Inference of macromolecular assemblies from crystalline state. J Mol Biol 372: 774–797.

38. Laskowski RA, Jabłońska J, Pravda L, Vařeková RS, Thornton JM (2018) PDBsum: Structural summaries of PDB entries. Protein Sci 27: 129–134.

39. Holm L, Rosenström P (2010) Dali server: conservation mapping in 3D. Nucleic Acids Res 38: W545–9.

40. Margraf T, Schenk G, Torda AE (2009) The SALAMI protein structure search server. Nucleic Acids Res 37: W480–4.

41. Unni S, Huang Y, Hanson RM, Tobias M, Krishnan S, Li WW et al. (2011) Web servers and services for electrostatics calculations with APBS and PDB2PQR. J Comput Chem 32: 1488–1491.

42. Pettersen EF, Goddard TD, Huang CC, Couch GS, Greenblatt DM, Meng EC et al. (2004) UCSF Chimera--a visualization system for exploratory research and analysis. J Comput Chem 25: 1605–1612.

43. Madeira F, Park YM, Lee J, Buso N, Gur T, Madhusoodanan N et al. (2019) The EMBL-EBI search and sequence analysis tools APIs in 2019. Nucleic Acids Res 47: W636–W641.

44. Waterhouse A, Bertoni M, Bienert S, Studer G, Tauriello G, Gumienny R et al. (2018) SWISS-MODEL: homology modelling of protein structures and complexes. Nucleic Acids Res 46: W296–W303.

45. Blanchet CE, Spilotros A, Schwemmer F, Graewert MA, Kikhney A, Jeffries CM et al. (2015) Versatile sample environments and automation for biological solution X-ray scattering experiments at the P12 beamline (PETRA III, DESY). J Appl Crystallogr 48: 431–443.

46. Franke D, Petoukhov MV, Konarev PV, Panjkovich A, Tuukkanen A, Mertens HDT et al. (2017) *ATSAS 2.8*: a comprehensive data analysis suite for small-angle scattering from macromolecular solutions. J Appl Crystallogr 50: 1212–1225.

47. Svergun DI (1999) Restoring low resolution structure of biological macromolecules from solution scattering using simulated annealing. Biophys J 76: 2879–2886.

48. Svergun DI, Petoukhov MV, Koch MH (2001) Determination of domain structure of proteins from X-ray solution scattering. Biophys J 80: 2946–2953.

49. Svergun DIBC, Barberato C, Koch MHJ (1995) CRYSOL–a program to evaluate X-ray solution scattering of biological macromolecules from atomic coordinates. Journal of applied crystallography 28: 768–773.

50. Altschul SF, Gish W, Miller W, Myers EW, Lipman DJ (1990) Basic local alignment search tool. J Mol Biol 215: 403–410.

51. Altschul SF, Madden TL, Schäffer AA, Zhang J, Zhang Z, Miller W et al. (1997) Gapped BLAST and PSI-BLAST: a new generation of protein database search programs. Nucleic Acids Res 25: 3389–3402.

52. Katoh K, Standley DM (2013) MAFFT multiple sequence alignment software version 7: improvements in performance and usability. Mol Biol Evol 30: 772–780.

53. Wong HC, Shin R, Krishna NR (2008) Solution structure of a double mutant of the carboxy-terminal dimerization domain of the HIV-1 capsid protein. Biochemistry 47: 2289–2297.

54. Byeon IJ, Meng X, Jung J, Zhao G, Yang R, Ahn J et al. (2009) Structural convergence between Cryo-EM and NMR reveals intersubunit interactions critical for HIV-1 capsid function. Cell 139: 780–790.

55. Cottee MA, Letham SC, Young GR, Stoye JP, Taylor IA (2020) Structure of Drosophila melanogaster ARC1 reveals a repurposed molecule with characteristics of retroviral Gag. Sci Adv 6: eaay6354.

56. Ma L, Jones CT, Groesch TD, Kuhn RJ, Post CB (2004) Solution structure of dengue virus capsid protein reveals another fold. Proc Natl Acad Sci U S A 101: 3414–3419.

57. Oliveira ERA, Mohana-Borges R, de Alencastro RB, Horta BAC (2017) The flavivirus capsid protein: Structure, function and perspectives towards drug design. Virus Res 227: 115–123.

58. Dokland T, Walsh M, Mackenzie JM, Khromykh AA, Ee KH, Wang S (2004) West Nile virus core protein; tetramer structure and ribbon formation. Structure 12: 1157–1163.

59. Ivanov D, Tsodikov OV, Kasanov J, Ellenberger T, Wagner G, Collins T (2007) Domain-swapped dimerization of the HIV-1 capsid C-terminal domain. Proc Natl Acad Sci U S A 104: 4353–4358.

60. Mattiroli F, Bhattacharyya S, Dyer PN, White AE, Sandman K, Burkhart BW et al. (2017) Structure of histone-based chromatin in Archaea. Science 357: 609–612.

61. Wang H, Wang M, Yang N, Xu RM (2015) Structure of the quaternary complex of histone H3-H4 heterodimer with chaperone ASF1 and the replicative helicase subunit MCM2. Protein Cell 6: 693–697.

62. Xie X, Kokubo T, Cohen SL, Mirza UA, Hoffmann A, Chait BT et al. (1996) Structural similarity between TAFs and the heterotetrameric core of the histone octamer. Nature 380: 316–322.

63. Stroud JC, Wu Y, Bates DL, Han A, Nowick K, Paabo S et al. (2006) Structure of the forkhead domain of FOXP2 bound to DNA. Structure 14: 159–166.

64. Moldoveanu T, Liu Q, Tocilj A, Watson M, Shore G, Gehring K (2006) The X-ray structure of a BAK homodimer reveals an inhibitory zinc binding site. Mol Cell 24: 677–688.

65. Brouwer JM, Westphal D, Dewson G, Robin AY, Uren RT, Bartolo R et al. (2014) Bak core and latch domains separate during activation, and freed core domains form symmetric homodimers. Mol Cell 55: 938–946.

66. Cowan AD, Smith NA, Sandow JJ, Kapp EA, Rustam YH, Murphy JM et al. (2020) BAK core dimers bind lipids and can be bridged by them. Nat Struct Mol Biol

67. Chang CK, Hou MH, Chang CF, Hsiao CD, Huang TH (2014) The SARS coronavirus nucleocapsid protein--forms and functions. Antiviral Res 103: 39–50.

68. Sandman K, Reeve JN (2006) Archaeal histones and the origin of the histone fold. Curr Opin Microbiol 9: 520–525.

69. Venkatesh S, Workman JL (2015) Histone exchange, chromatin structure and the regulation of transcription. Nat Rev Mol Cell Biol 16: 178–189.

70. Chu YP, Chang CH, Shiu JH, Chang YT, Chen CY, Chuang WJ (2011) Solution structure and backbone dynamics of the DNA-binding domain of FOXP1: insight into its domain swapping and DNA binding. Protein Sci 20: 908–924.

71. Volle C, Dalal Y (2014) Histone variants: the tricksters of the chromatin world. Curr Opin Genet Dev 25: 8–14, 138.

72. Colpitts TM, Barthel S, Wang P, Fikrig E (2011) Dengue virus capsid protein binds core histones and inhibits nucleosome formation in human liver cells. PLoS One 6: e24365.

73. Salery M, Dos Santos M, Saint-Jour E, Moumné L, Pagès C, Kappès V et al. (2017) Activity-Regulated Cytoskeleton-Associated Protein Accumulates in the Nucleus in Response to Cocaine and Acts as a Brake on Chromatin Remodeling and Long-Term Behavioral Alterations. Biol Psychiatry 81: 573–584.

74. Czabotar PE, Westphal D, Dewson G, Ma S, Hockings C, Fairlie WD et al. (2013) Bax crystal structures reveal how BH3 domains activate Bax and nucleate its oligomerization to induce apoptosis. Cell 152: 519–531.

75. Sibanda BL, Thornton JM (1985) Beta-hairpin families in globular proteins. Nature 316: 170–174.

76. Duddy WJ, Nissink JW, Allen FH, Milner-White EJ (2004) Mimicry by asx- and ST-turns of the four main types of beta-turn in proteins. Protein Sci 13: 3051–3055.

77. Ivanov D, Stone JR, Maki JL, Collins T, Wagner G (2005) Mammalian SCAN domain dimer is a domain-swapped homolog of the HIV capsid C-terminal domain. Mol Cell 17: 137–143.

78. Zhao G, Perilla JR, Yufenyuy EL, Meng X, Chen B, Ning J et al. (2013) Mature HIV-1 capsid structure by cryo-electron microscopy and all-atom molecular dynamics. Nature 497: 643–646.

79. Worthylake DK, Wang H, Yoo S, Sundquist WI, Hill CP (1999) Structures of the HIV-1 capsid protein dimerization domain at 2.6 A resolution. Acta Crystallogr D Biol Crystallogr 55: 85–92.

80. Obal G, Trajtenberg F, Carrión F, Tomé L, Larrieux N, Zhang X et al. (2015) STRUCTURAL VIROLOGY. Conformational plasticity of a native retroviral capsid revealed by x-ray crystallography. Science 349: 95–98.

81. Bailey GD, Hyun JK, Mitra AK, Kingston RL (2009) Proton-linked dimerization of a retroviral capsid protein initiates capsid assembly. Structure 17: 737–748.

82. Dodonova SO, Prinz S, Bilanchone V, Sandmeyer S, Briggs JAG (2019) Structure of the Ty3/Gypsy retrotransposon capsid and the evolution of retroviruses. Proc Natl Acad Sci U S A 116: 10048–10057.

83. Nielsen LD, Pedersen CP, Erlendsson S, Teilum K (2019) The Capsid Domain of Arc Changes Its Oligomerization Propensity through Direct Interaction with the NMDA Receptor. Structure 27: 1071–1081.e5.

84. Alfadhli A, Steel E, Finlay L, Bächinger HP, Barklis E (2002) Hantavirus nucleocapsid protein coiled-coil domains. J Biol Chem 277: 27103–27108.

85. Kammerer RA, Schulthess T, Landwehr R, Lustig A, Engel J, Aebi U et al. (1998) An autonomous folding unit mediates the assembly of two-stranded coiled coils. Proc Natl Acad Sci U S A 95: 13419–13424.

86. Steinmetz MO, Jelesarov I, Matousek WM, Honnappa S, Jahnke W, Missimer JH et al. (2007) Molecular basis of coiled-coil formation. Proc Natl Acad Sci U S A 104: 7062–7067.

87. Bhargav SP, Vahokoski J, Kallio JP, Torda AE, Kursula P, Kursula I (2015) Two independently folding units of Plasmodium profilin suggest evolution via gene fusion. Cell Mol Life Sci 72: 4193–4203.

88. Han H, Kursula P (2014) Periaxin and AHNAK nucleoprotein 2 form intertwined homodimers through domain swapping. J Biol Chem 289: 14121–14131.

89. Smyth DR, Kalitsis P, Joseph JL, Sentry JW (1989) Plant retrotransposon from Lilium henryi is related to Ty3 of yeast and the gypsy group of Drosophila. Proc Natl Acad Sci U S A 86: 5015–5019.

