## Supplementary material for "Crystal and solution structures reveal oligomerization of individual capsid homology domains of Drosophila Arc": S1 table

S1 Table. The expression constructs used in the current study.

| Construct | Uniprot entry | Residues | N-terminal extension |
| --- | --- | --- | --- |
| dArc1-NL | Q7K1U0 | 51-122 | SGSG |
| dArc1-CL | Q7K1U0 | 123-208 | SGSG |
| dArc2-NL | Q7JV70 | 38-109 | S |
| dArc2-CL | Q7JV70 | 110-193 | SGSG |
| hArc-NL | Q7LC44 | 207-277 | GAMG |
| hArc-CL | Q7LC44 | 277-370 | GAM |
