## Supplementary material for "Crystal and solution structures reveal oligomerization of individual capsid homology domains of Drosophila Arc": Fig S1

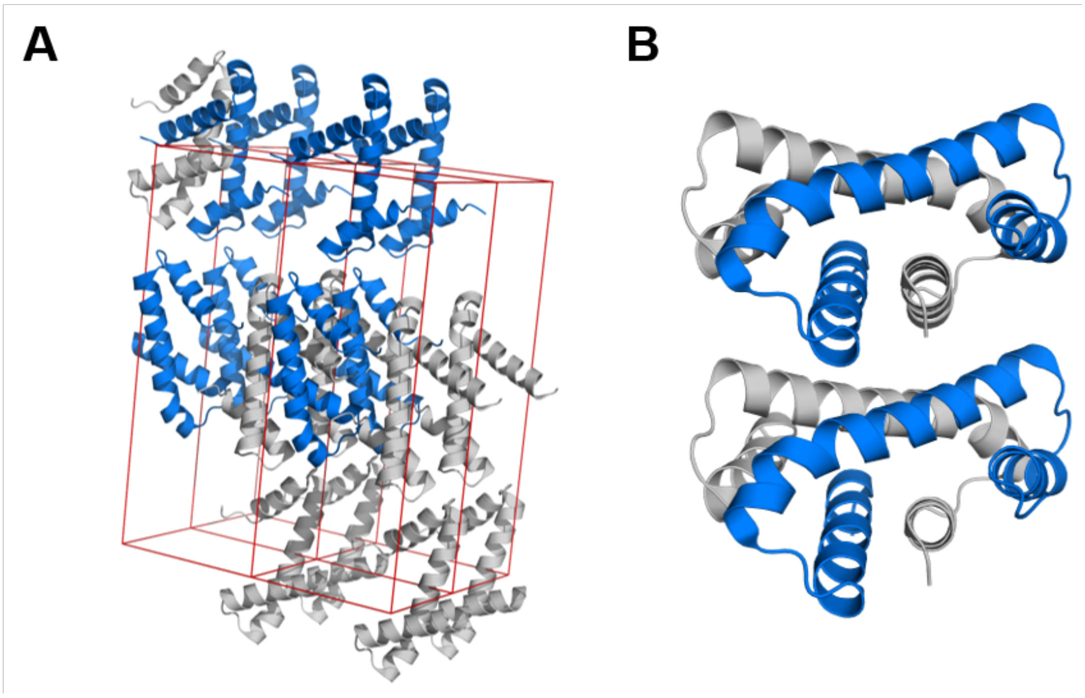


**Fig. S1. Crystal packing in dArc2-NL.** (A) The unit cell. (B) Stacking of dArc2-NL domain-swapped dimers in the lattice.
