## Supplementary figures and images for "Crystal and solution structures reveal oligomerization of individual capsid homology domains of Drosophila Arc"

### Fig S2

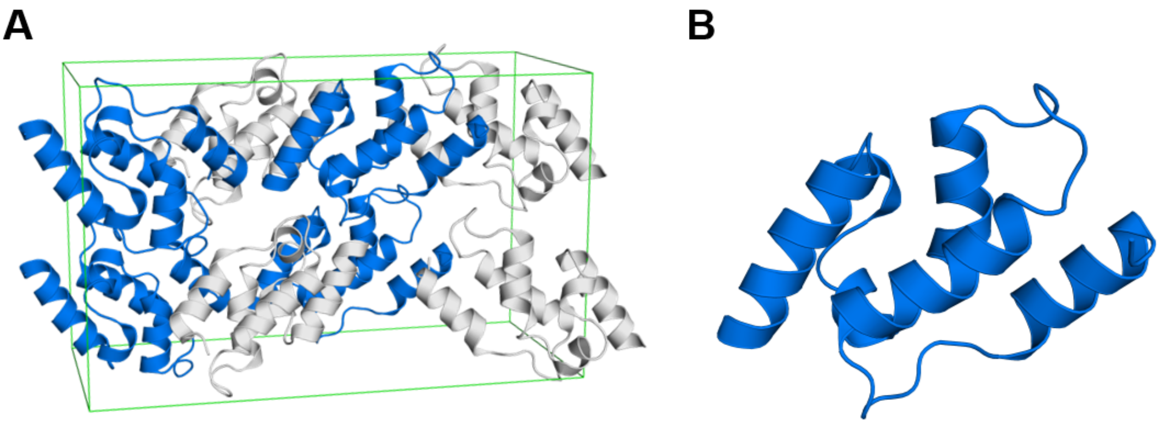


**Fig. S2.** Crystal packing in dArc1-CL. (A) The unit cell. (B) A monomer of dArc1-CL.
