## Supplementary material for "Crystal and solution structures reveal oligomerization of individual capsid homology domains of Drosophila Arc": Fig S3

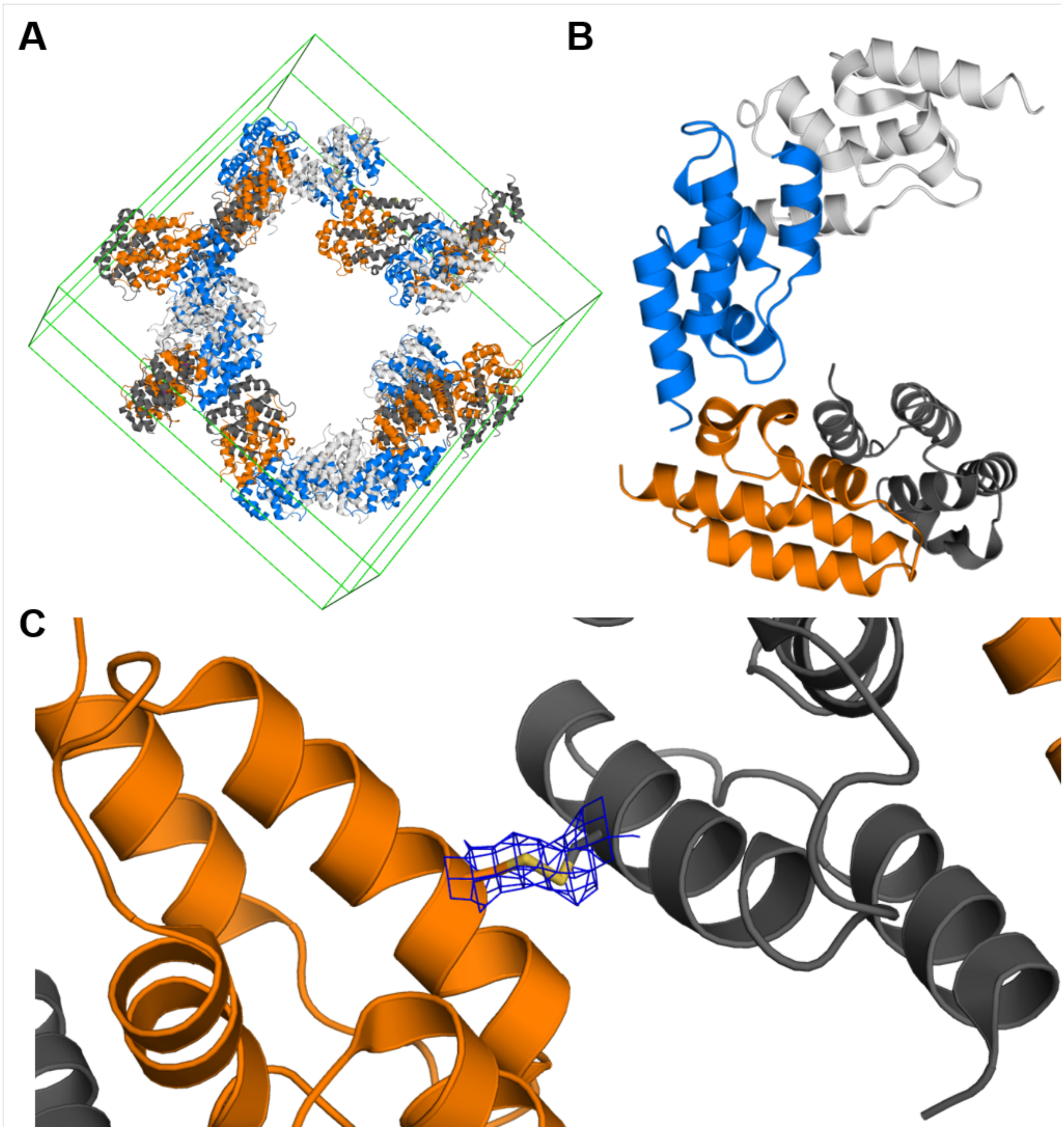


**Fig. S3.** **Crystal packing in dArc2-CL.** (A) The unit cell. (B) dArc2-CL is packed as homodimers. (C) In the crystal lattice, dimers are linked to each other *via* a disuplhide bridge.
